# Distinct Stromal Cell Populations Define the B-cell Acute Lymphoblastic Leukemia Microenvironment

**DOI:** 10.1101/2024.09.10.612346

**Authors:** Mauricio N. Ferrao Blanco, Bexultan Kazybay, Mirjam Belderbos, Olaf Heidenreich, Hermann Josef Vormoor

## Abstract

The bone marrow microenvironment plays a critical role in B-cell acute lymphoblastic leukemia (B-ALL) progression, yet its cellular heterogeneity remains poorly understood. Using single-cell RNA sequencing on patient-derived of bone marrow aspirates from pediatric B-ALL patients, we identified two distinct mesenchymal stromal cell (MSC) populations: early mesenchymal progenitors and adipogenic progenitors. Spatial transcriptomic analysis further revealed the localization of these cell types and identified a third stromal population, osteogenic-lineage cells, exclusively present in the bone biopsy. Functional *ex vivo* assays using sorted stromal populations derived from B-ALL patient bone marrow aspirates demonstrated that both early mesenchymal and adipogenic progenitors secrete key niche-supportive factors, including CXCL12 and Osteopontin, and support leukemic cell survival and chemoresistance. Transcriptomic profiling revealed that B-ALL cells interact differently with stromal subtypes. Notably, adipogenic progenitors, but not early mesenchymal progenitors, provide support to leukemic cells through interleukin-7 and VCAM1 signaling. Stromal cells from B-ALL patients exhibited an enhanced adipogenic differentiation capacity compared to healthy controls. Moreover, co-culture experiments showed that B-ALL cells induce adipogenic differentiation in healthy MSCs through a cell contact-dependent mechanism. Adipogenic progenitors were also enriched in relapse samples, implicating them in disease progression. These findings highlight the complexity of the B-ALL microenvironment and identify different specialized stromal niches with which the leukemic cells can engage.

## Introduction

The bone marrow microenvironment is a highly specialized tissue consisting of different cell types and extracellular matrix proteins, essential for regulating hematopoiesis and skeletal regeneration [1]. Central to this intricate network are mesenchymal stromal cells (MSCs), a heterogeneous population encompassing multipotent progenitors of skeletal lineages [2, 3]. These MSCs contribute to the formation of specialized niches within the bone marrow, which provide essential support for hematopoietic stem cell (HSC) function, including homing, quiescence, and lineage commitment [4–6].

The dynamic interplay between leukemic cells and the bone marrow microenvironment is increasingly recognized as a critical factor in hematological malignancies, including B-cell acute lymphoblastic leukemia (B-ALL) [7]. Leukemic cells actively reprogram their microenvironment to create a supportive niche that promotes survival, proliferation, and resistance to therapy [8–10]. While previous studies have implicated a key role of the bone marrow microenvironment in B-ALL, a comprehensive understanding of its cellular heterogeneity remains elusive.

To address this knowledge gap, we conducted a comprehensive single-cell RNA sequencing analysis of the bone marrow microenvironment in pediatric B-ALL. By systematically characterizing the stromal, hematopoietic, and leukemic compartments, we identified distinct MSC populations as key contributors to the leukemia-supportive niche. Notably, our findings reveal a previously unrecognized role for adipogenic progenitors in the leukemic niche and shed light on the complexity of the cellular micro-environment in B-ALL.

## Results

### The single cell bone marrow landscape of B-ALL

To comprehensively characterize the bone marrow microenvironment in B-ALL, we employed flow cytometry to isolate leukemic plus normal B-lineage (CD19+), non-leukemic hematopoietic (CD45+CD235a+), and non-hematopoietic (CD19-CD45-CD235a-) mononuclear cells from patient bone marrow aspirates (Figure 1A). The identity of the non-hematopoietic fraction was confirmed by its ability to form colony-forming unit fibroblasts (CFU-Fs), express MSC markers (CD90, CD73, CD105), and differentiate into adipocytes and osteoblasts (Supplementary Figure 1). We analyzed bone marrow mononuclear cells from nine B-ALL patients, encompassing 5 different molecular subtypes as well as samples taken at diagnosis and relapse (Figure 1B; Supplementary Table 1). A healthy donor sample from an 18-year-old woman was included in the scRNA-seq analysis and only the non-hematopoietic compartment from this sample was sorted and sequenced. After quality control and doublet exclusion, 28,333 high-quality cells were obtained and integrated into a single dataset for further analysis. We generated a single-cell transcriptomic map of the integrated stromal, hematopoietic, and leukemic compartment (Figure 1C). We observe variable contribution of each patient to each cluster (Figure 1D), though given our small cohort (n=9), an association with molecular subtype, age and sex is not feasible. Unsupervised clustering identified twenty major cell populations based on the expression of lineage-specific markers (Figure 1E; Supplementary Table 2). These clusters included, among others, stromal cells (expressing *CXCL12*, *THY1* and *FN1*), mature B cells (expressing *CD19*, *CD22* and *MS4A1*) and leukemic cells (expressing *CD19*, *CD22* and *MME*). The expression of CD19 was relatively low, which could be associated to technical limitations of scRNAseq, such as high proportion of zero counts resulting from low sequencing depth [11], the lower stability of mRNA compared to protein, or biological factors including acquired mutations or alternative splicing [12, 13]. Expression of the hematopoietic marker *PTPRC,* which encodes CD45, was ubiquitous except in erythroblasts and stromal cells. Further discrimination between leukemic and mature B cells was based on the aberrant karyotype of ALL cells, inferred by copy number variation analysis (Supplementary Figure 2). Unbiased cell type recognition using bone marrow scRNAseq references [14, 15], confirmed the cell annotation and mapped leukemic cells to pro-B and preproB-cells, confirming their immature phenotype (Supplementary Figure 3). Gene ontology enrichment analysis validated the cellular identity of each cluster, for example, stromal cells were enriched in genes associated with collagen-containing extracellular matrix, while T cells were enriched in express genes associated with T cell activation (Supplementary Figure 4).

**Figure 1.**
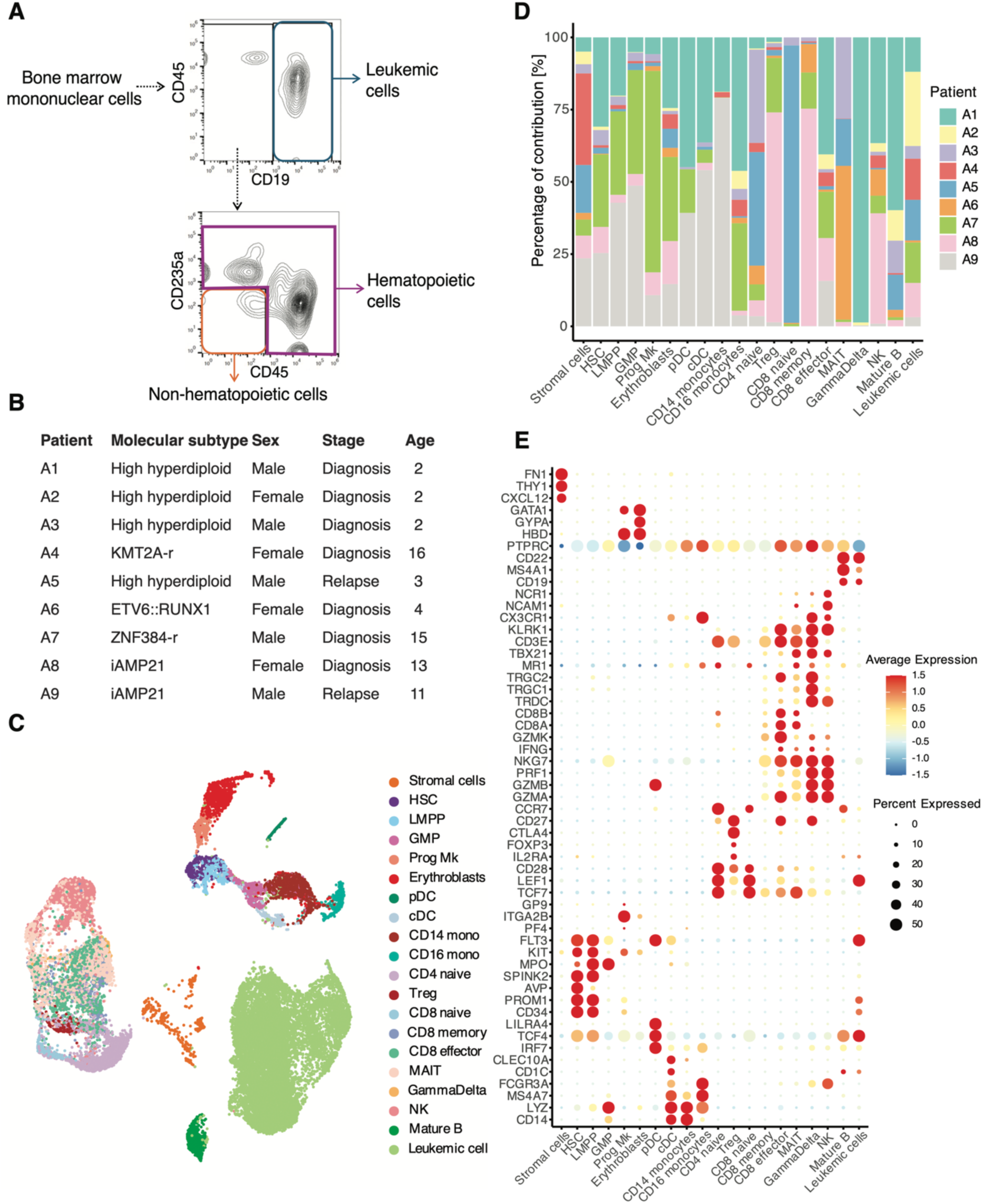
Integrated analysis of the B-ALL bone marrow microenvironment. A) Gating strategy for the enrichment of leukemic cells plus normal B-lineage cells (CD19+), hematopoietic cells (CD45+CD235a+) and non-hematopoietic cells (negative for all markers). B) Table detailing the molecular subtype, age and sex associated with each sample sequenced. C) Uniform Manifold Approximation and Projection (UMAP) plot showing cell clusters from the analysis of 28,333 cells integrated from 9 patient samples. D) Barplot indicating the contribution of individual patients to each cell cluster. E) DotPlot representation of gene expression of typical cell cluster markers, 3 markers per cluster. The size of the dot indicates the percentage of cells expressing the gene and the color indicates the average expression level. Abbreviations. Lymphoid-primed multipotent progenitors (LMPP), megakaryocyte progenitors (Prog Mk), granulocyte-macrophage progenitors (GMP), plasmacytoid dendritic cells (pDC), conventional dendritic cells (cDC), hematopoietic stem cells (HSCs), mucosal-associated invariant T cells (MAIT), regulatory T cells (Tregs), natural killer cells (NK).

In summary, the data presented here provide a comprehensive single-cell transcriptomic blueprint of the bone marrow microenvironment in pediatric B-ALL.

### Identification of distinct stromal cell populations

The stromal landscape in B-ALL remains largely unexplored. In-depth analysis of the stromal cell cluster revealed two distinct populations: early mesenchymal progenitors and adipogenic progenitors (Figures 2A). By comparing our data with a human bone marrow stromal cell atlas [14], we mapped early mesenchymal progenitors to fibro-MSCs (Figure 2B and Supplementary Figure 5). In contrast, adipogenic progenitors mapped to adipo-MSCs, which resemble reported marrow adipogenic lineage precursors and Adipo-CXCL12-abundant-reticular (Adipo-CAR) cells in mice [16, 17]. Accordingly, adipogenic progenitors showed high expression of adipogenic markers *LPL, PPARG* and *LEPR*, while early mesenchymal progenitors did not, neither markers of other stromal fate-committed cells such as osteoblasts (*IBSP, RUNX2, BGLAP*) or chondrocytes (*SOX9*) (Figure 2D). Gene set enrichment analysis demonstrated that adipogenic progenitors were enriched in extracellular matrix-related genes (Figure 2C), mainly collagen type I, type III and type XII, as well as fibrillin-1 and Decorin (Figure 2D).

**Figure 2.**
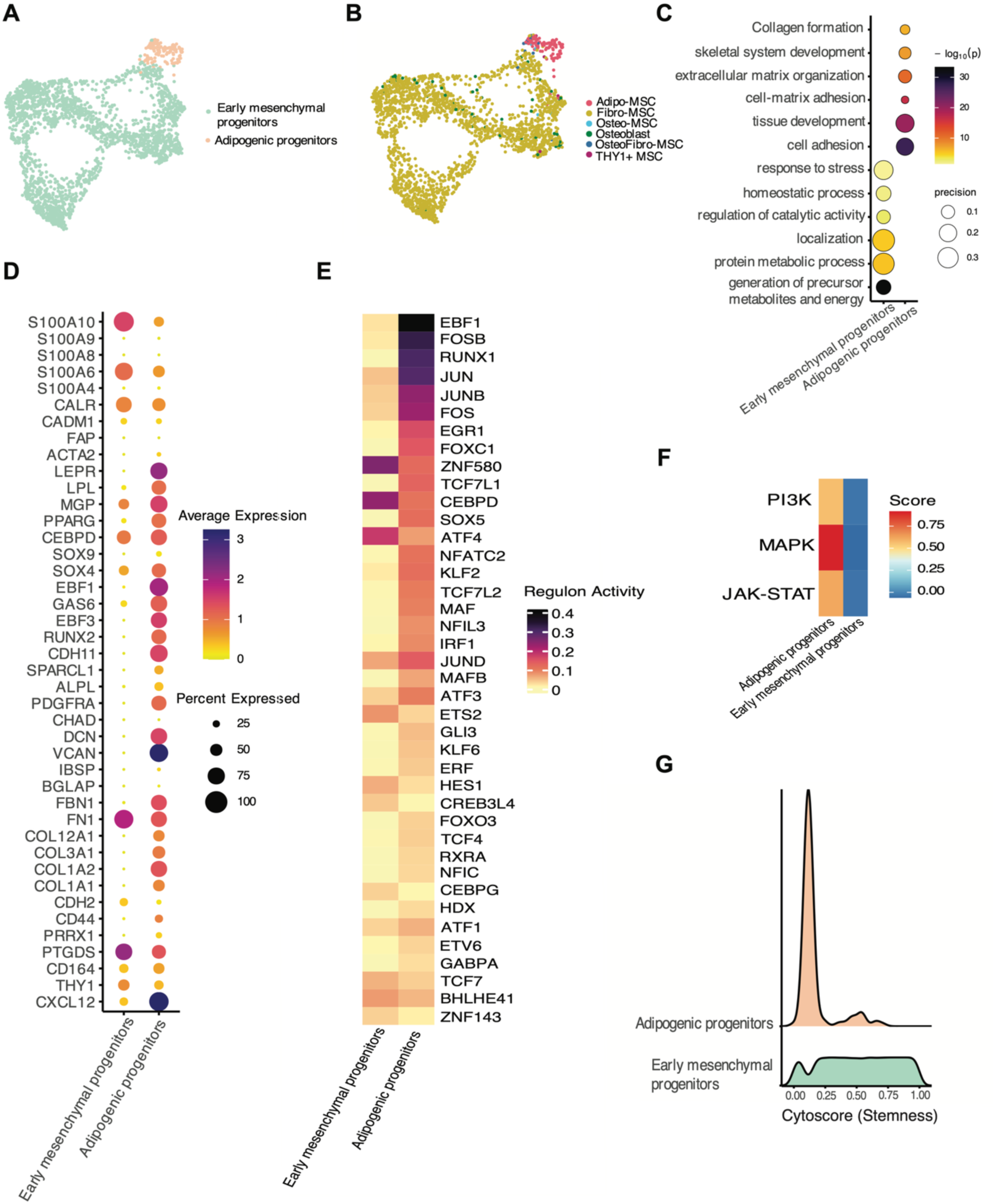
Characterization of the B-ALL stromal niche. A) UMAP representation of the stromal clusters. B) UMAP plot showing the annotation after mapping the stromal cluster to the reported by Bandyopadhyay et al, 2024. C) Dot plot showing enriched gene ontology (GO) terms of differentially expressed genes in the cell clusters. D) Dot plot representing the gene expression of selected markers in adipogenic and early mesenchymal progenitors. E) Heatmap representation of transcription factor activity score inferred by SCENIC. F) Heatmap depicting pathway activity analysis. G) Ridge plot showing the distribution of Cytotrace scores for adipogenic and early mesenchymal progenitors. Each ridge represents a different stromal cell population, with adipogenic progenitors in pink and early mesenchymal progenitors in green. The x-axis represents the Cytotrace score, which reflects the differentiation trajectory of cells. A higher Cytotrace score indicates a higher stemness profile. The y-axis represents the density of cells at a particular Cytotrace score.

To elucidate transcription factor activity of stromal cells, we assessed the activity of key transcription factors using the single-cell regulatory network inference and clustering (SCENIC) method [18]. Adipogenic progenitors exhibited high activity of adipogenic master regulators such as EBF1 and FOSB [19, 20], while early mesenchymal progenitors showed elevated ZNF580, CEBPD and ATF4 activity, associated with a more upstream progenitor state [21] (Figure 2E, Supplementary Table 3). Signaling pathway analysis suggest higher MAPK, PI3K and JAK/STAT signaling in adipogenic progenitors compared with early mesenchymal progenitors, consistent with their role in adipogenesis [22–24] (Figure 2F). Lastly, pseudotime analysis indicated a more primitive state for early mesenchymal progenitors compared to adipogenic progenitors (Figure 2G).

Overall, our findings delineate the transcriptomic diversity within the stromal niche in B-ALL, providing novel insights into the cellular heterogeneity of this critical microenvironment.

### Spatial localization of stromal niches

We next analyzed a publicly available spatial transcriptomics dataset generated using a trephine bone biopsy sample from a B-ALL patient (Xenium technology, 10x Genomics) [25]. Patient age, sex, or molecular subtype were not provided with the dataset. Due to the limitation of the Xenium gene panel, which includes only 477 genes, a detailed annotation of hematopoietic clusters-similar to that achieved in our single-cell RNA-seq analysis of bone marrow aspirates-was not feasible (Supplementary Figure 6). To investigate the spatial localization of stromal populations in B-ALL bone marrow, we annotated distinct stromal cell types, identifying early mesenchymal progenitors, adipogenic progenitors, and osteogenic-lineage cells within the sample (Figure 3A). Cell types were identified based on the expression of top marker genes: adipogenic progenitors expressed high levels of LPL and FABP4, osteogenic-lineage cells expressed BMP2 and IBSP, while early mesenchymal progenitors lacked lineage-committed markers but expressed the extracellular matrix gene COL1A1 (Figure 3B). The stromal niche was predominantly composed of adipogenic progenitors, which accounted for approximately 70% of the stromal cells (Figure 3C). Notably, osteogenic-lineage cells-absent in our single-cell RNA seq dataset of bone marrow aspirates-were identified in this spatial dataset, likely due to their strong adherence to the bone surface, making them less accessible in aspirated material.

**Figure 3.**
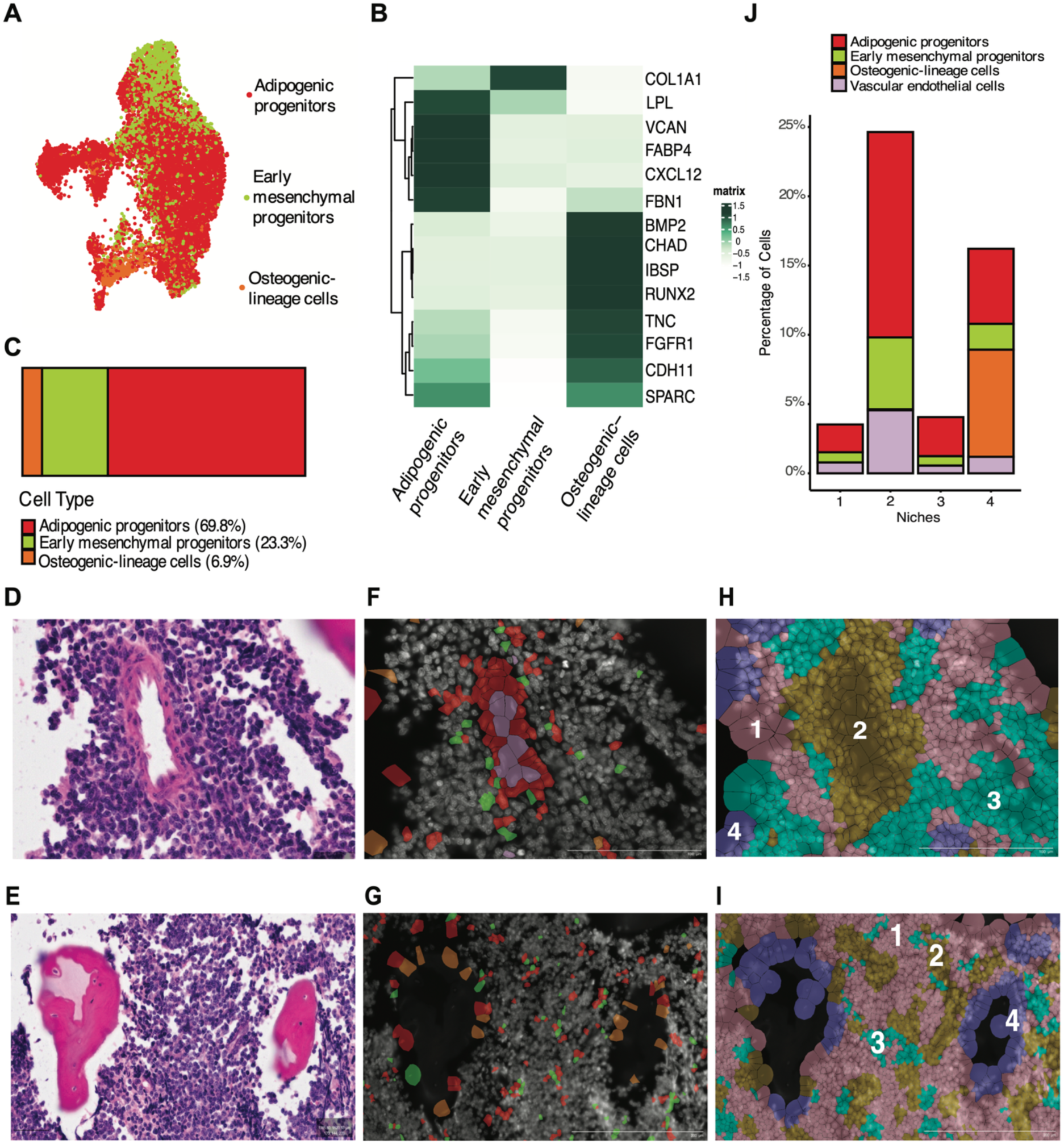
Spatial single-cell analysis of the B-ALL bone marrow niche. A) Uniform Manifold Approximation and Projection (UMAP) plot showing the stromal cell clusters identified in the Xenium data. B) Heatmap showing the average gene expression per cluster of selected top markers. C) Barplot representing the percentage of each stromal population relative to the total stromal cell types in the dataset. D-E) Hematoxilin & Eosin staining of the bone biopsy, zooming in a blood vessel (D) and trabelular bone (E). F-G) Localization of early mesenchymal progenitors (green), adipogenic progenitors (red), osteogenic-lineage cells (brown) and vascular endothelial cells (violet) on the same area as in D and E. H-I) Localization of the 4 different niches on the same area as in D and E. Each color represents a different niche, pink (1), yellow (2), cyan (3), purple (4). J) Barplot showing the cell type distribution between the four niches. In the y-axis, the percentage of these cell types relative to all the cell types composing the niche is shown.

Interestingly, adipogenic progenitors were primarily located around blood vessels and closely associated with vascular endothelial cells (Figure 3D & F). This spatial pattern suggests that at least a subset of these cells may represent pericytes, which is consistent with previous murine studies linking adipogenic progenitors to perivascular cells [16]. In contrast, early mesenchymal progenitors were more randomly distributed throughout the marrow, while osteogenic-lineage cells were predominantly located along the bone surface (Figure 3E & G).

A niche analysis incorporating all cell populations within the spatial transcriptomic dataset revealed distinct spatial domains, underscoring the differential localization of stromal populations within the B-ALL bone marrow microenvironment (Figure 3J; Supplementary Figure 7). Osteogenic-lineage cells were exclusively found in **niche 4**, whereas adipogenic progenitors and early mesenchymal progenitors were mainly enriched in **niche 2**, together with vascular endothelial cells (Figure 3H & I).

These findings emphasize the spatial heterogeneity and distinct localization patterns of stromal populations within the bone marrow ecosystem in B-ALL.

### Functional heterogeneity of mesenchymal stromal cell populations

To further validate and functionally characterize the distinct stromal populations observed in bone marrow aspirates, we developed a FACS sorting strategy based on markers identified through our scRNA-seq analysis of non-cultured stromal cells (Figure 4A & 4B). Adipogenic progenitors were defined by the high expression of *CDH11* (Cadherin 11) and *VCAM1* (CD106), while early mesenchymal progenitors were negative for these markers and positive for CD81 and *THY* (CD90). Notably, commonly used MSC markers exhibited variable expression among stromal subsets (Figure 4A). Flow cytometry analysis of cultured, sorted stromal populations confirmed the expression patterns of these markers (Figure 4H). Cadherin 11 and CD106 were detected on the surface of adipogenic progenitors (100% and 30%, respectively), while early mesenchymal progenitors were negative for these markers. Conversely, CD90 was consistently detected on early mesenchymal progenitors (99.5%) but was only partially expressed on adipogenic progenitors (40%). CD81 was expressed on both populations (100%). Both populations also exhibited strong positivity for the stromal markers CD51 and CD164 (>98% in both). These results highlight the importance of using a comprehensive panel of markers when studying MSCs and underscore the distinct phenotypic characteristics of these two stromal populations.

**Figure 4.**
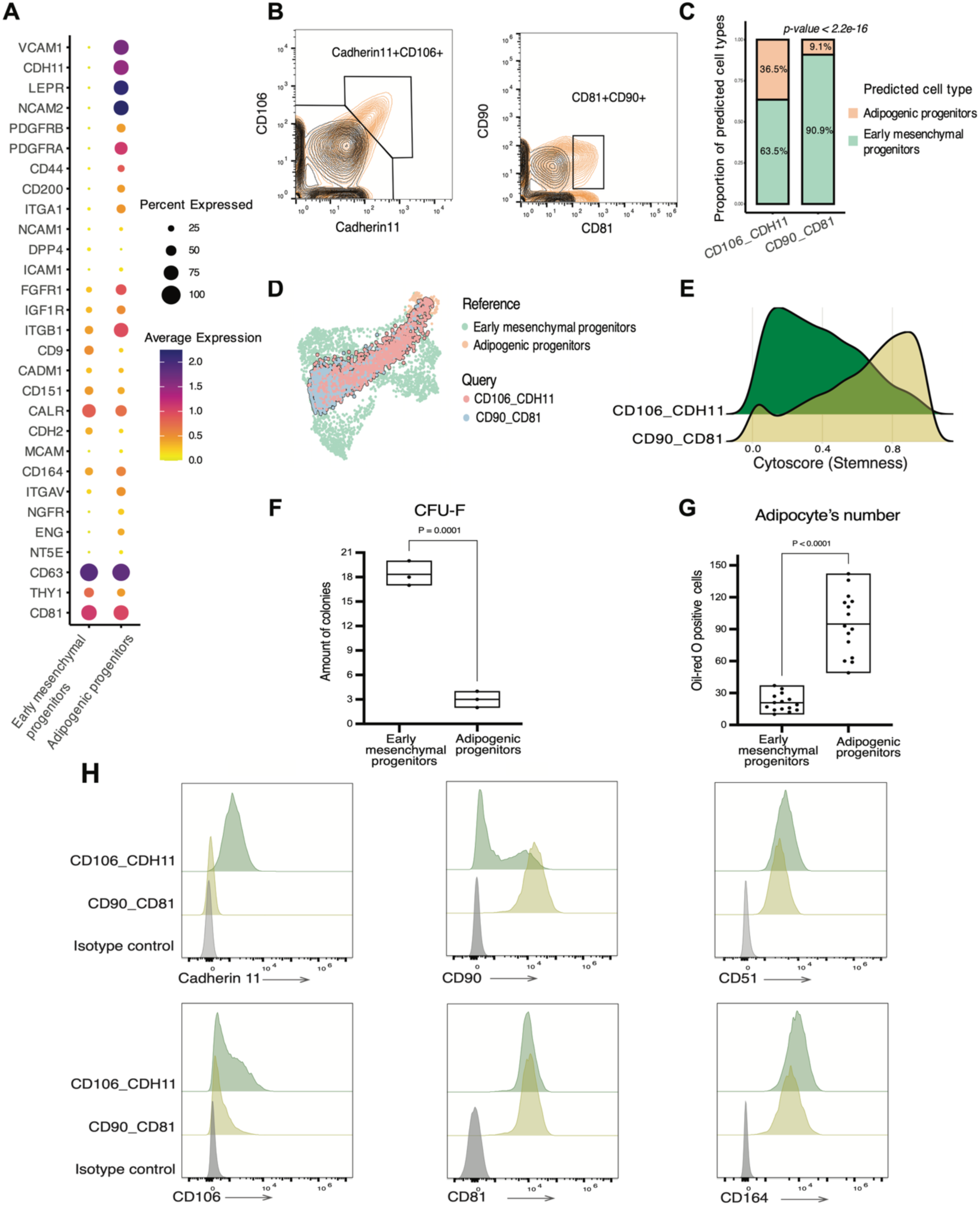
Functional characterization of stromal cell populations in B-ALL. A) Dotplot representation of selected cell surface markers in non-cultured early mesenchymal and adipogenic progenitors derived from bone marrow aspirates. The size of the dot indicates the percentage of cells expressing the gene and the color indicates the average expression level. B) Gating strategy for the sorting of stromal populations. In black the isotype control, while in orange the stained sample. C) Barplot showing the reference-based annotation of the sorted cultured stromal cells, using the non-expanded stromal cells as a reference. Sorted stromal cells were analyzed at passage 3. D) UMAP representation of the scRNAseq data of the two cultured stromal populations projected in the reference dataset of non-expanded stromal cells. E) Ridge plots of the two sorted stromal populations showing the predicted order score in differentiation. Each ridge represents a different stromal cell population, with CD106+CDH11+ in green and CD81+CD90+ in yellow. The x-axis represents the Cytotrace score, which reflects the differentiation trajectory of cells. A higher Cytotrace score indicates a higher stemness profile. The y-axis represents the density of cells at a particular Cytotrace score. F) CFU-F analysis of sorted stromal populations cultured *ex vivo* at passage 3. A total of 475 cells per well were plated in a 6-well plate (50 cells/cm2). Colonies were quantified after 10 days of culture (n=3). Statistical significance was assessed by a two-tailed unpaired Student’s t-test. G) Stromal cells at passage 3 were cultured in adipogenic medium for 14 days. Oil red O stains lipid droplets (red). The number of positive cells were quantified per well in a 96-well plate. Data is presented as mean ± standard deviation. Statistical significance was assessed by a two-tailed unpaired Student’s t-test. H) Immunophenotypic characterization of cultured, sorted stromal populations at passage 3. Histograms showing the fluorescence intensity of CD markers determined by flow cytometry. Signal intensity is shown on the x-axis and count on the y-axis. Adipogenic progenitors are shown in green, early mesenchymal progenitors in yellow and isotype controls in gray.

To validate our sorting strategy, we performed single-cell RNA sequencing on sorted early mesenchymal and adipogenic progenitors and assessed their similarity to the sorted – non cultured stromal populations signature (Figure 4C & D). By annotating the cultured stromal populations using uncultured ones as a reference, we observed that 36.5% of the CD106+CDH11+ cells were adipogenic progenitors in contrast to only 9.1% in case of the CD81+CD90+ cells, confirming the sorting strategy for the stromal population enrichment. Pseudotime analysis confirmed the more differentiated state of adipogenic progenitors compared to early mesenchymal progenitors (Figure 4E). Functional assays revealed a higher CFU-F formation capacity and proliferation rate in early mesenchymal progenitors, consistent with their stem/progenitor-like phenotype (Figure 4F). Conversely, adipogenic progenitors demonstrated enhanced adipogenic and osteogenic differentiation potential (Figures 4G, Supplementary Figure 8). Overall, these findings confirm the functional divergence of early mesenchymal and adipogenic progenitors.

### Early mesenchymal and adipogenic progenitors support leukemic cell survival via alternative mechanisms

Leukemic cell survival is dependent on the microenvironment [26]. To investigate whether stromal heterogeneity influences the production of different supportive factors, we analyzed the expression of known niche factors [26, 27]. RNA expression indicated that different MSC types may produce specific support factors; for example, *SPP1* (osteopontin) and CDH2 (N-Cadherin) was highest in early mesenchymal progenitors, whereas *CXCL12*, *VCAM1* (vascular cell adhesion molecule-1), *ANGPT1* (angiopoietin-1), *IL7* (interleukin 7) and *KITLG* (stem cell factor) were predominantly expressed by adipogenic progenitors (Figure 5A) [28] [29] [30]. To further explore stromal-leukemic cell interactions in an unbiased manner, we performed ligand-receptor interaction analysis (Figure 5B). By quantifying the interactions of the niche cells with the leukemic cells, we found that adipogenic progenitors were categorized as the primary source of ligands interacting with leukemic cells, followed by early mesenchymal progenitors (Figure 5C).

**Figure 5.**
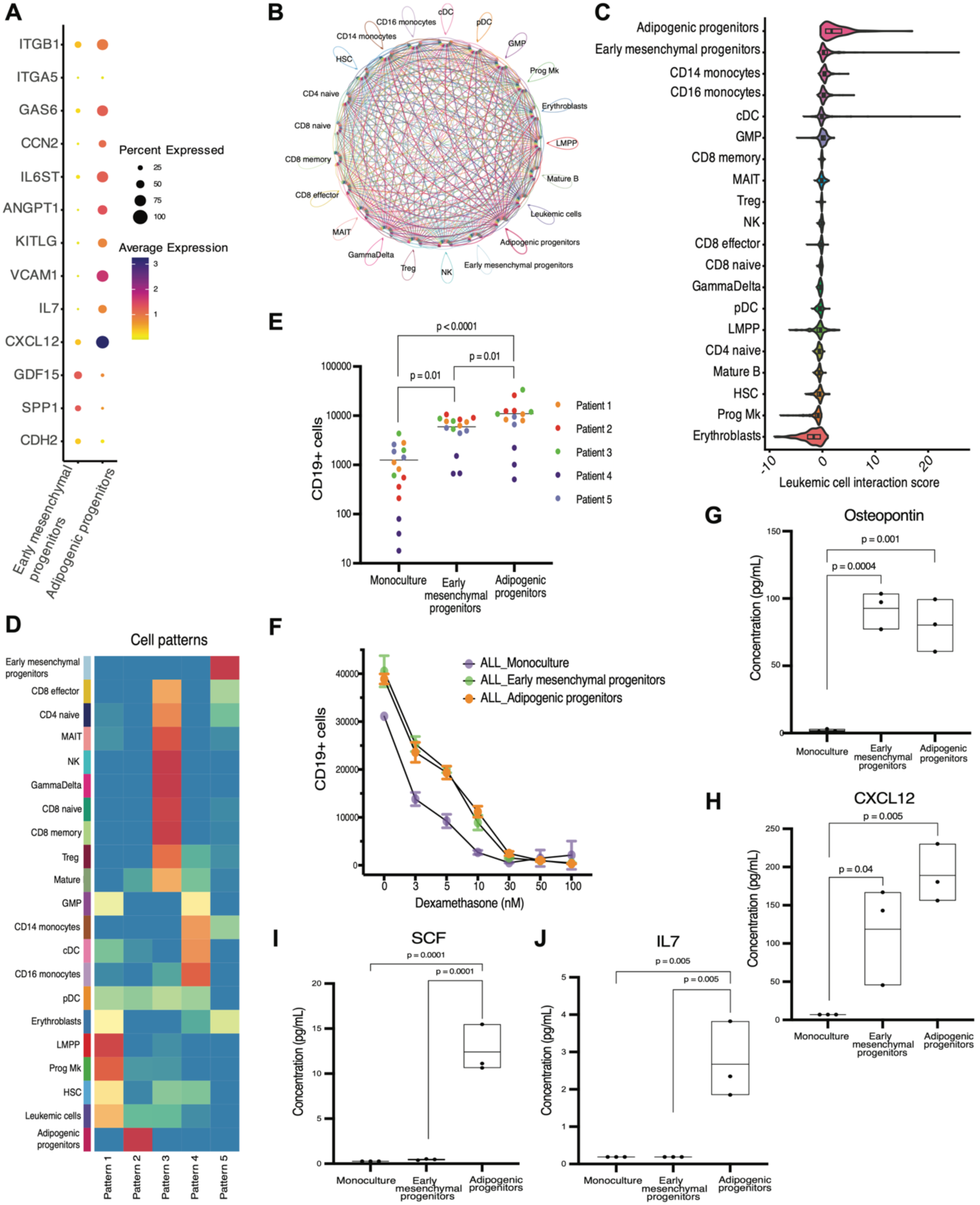
Cell-cell communication analysis in the B-ALL niche. A) Dotplot representation of the gene expression of widely studied niche factors. The size of the dot indicates the percentage of cells expressing the gene and the color indicates the average expression level. B) Circle plot showing the interactions between niche cell clusters and leukemic cells. The thickness of the line represents the strength of the interaction. The colors of the lines represent a different cell population. C) Representation of the strength of the cell-cell interaction of the niche cells with the leukemic cells, calculated as a leukemic cell interaction score. The score is based on the average expression of signaling factors (cytokines, integrins, extracellular matrix components) that were inferred to interact with leukemic cells. D) Non-negative matrix factorization (NMF) to identify modules in the cell-cell interaction pathways between niche cells and leukemic cells. A total of 5 modules were identified based on Cophenetic and Silhouette metrics. Clusters of cells are grouped into each module. E) Absolute number of CD19+ leukemic cells per well after seven days of monoculture or co-culture with adipogenic and early mesenchymal progenitors. Colors depict different patients. Data is presented as mean ± standard deviation. Statistical significance was assessed by a two-way ANOVA with Bonferroni’s multiple comparison. F) Dose-response analysis of Dexamethasone (0 – 100 nM) by quantifying the absolute number of CD19+ leukemic cells per well after seven days of monoculture or co-culture with adipogenic and early mesenchymal progenitors. Cell counts are shown as mean ± standard deviation of three replicates. A two-way ANOVA revealed significant effects of Dex concentration (**p** < 2 × 10⁻¹⁶), cell type (**p** < 2 × 10⁻¹⁶), and their interaction (**p** = 9.81 × 10⁻¹⁵). Pairwise comparisons (adjusted using the Benjamini-Hochberg method) at each Dex concentration revealed significant differences between groups: 3 μM Monoculture vs. AP (p = 0.015), EMP vs. AP (p = 0.035), 5 μM Monoculture vs. EMP (p = 6.8 × 10⁻⁵), Monoculture vs. AP (p = 6.8 × 10⁻⁵), 10 μM Monoculture vs. EMP (p = 0.00081), Monoculture vs. AP (p = 0.00029), 30 μM Monoculture vs. EMP (p = 0.0077), Monoculture vs. AP (p = 0.00056), EMP vs. AP (p = 0.0077). G-J) Multiplex cytokine analysis of osteopontin, CXCL12, SCF and IL-7 in the medium of the co-culture experiments. Each datapoint denotes a replicate sample. Statistical significance was assessed by a one-way ANOVA and Tukey’s HSD post-hoc test.

To gain a deeper understanding of the underlaying regulatory mechanisms in the complex communication ecosystem, we further classified the cell-cell interactions into patterns where groups of cells are grouped together if they share a similar communication pattern (Figure 5D). We identified 5 patterns, each of them related to a group of cells, hematopoietic progenitors (pattern 1), adipogenic progenitors (pattern 2), T lymphocytes (pattern 3), myeloid cells (pattern 4) and early mesenchymal progenitors (pattern 5). These patterns were characterized by different interactions involving secreted factors, cell-cell contact, and extracellular matrix-receptor signaling, highlighting the different mechanisms that cells use in the bone marrow niche (Supplementary Figure 9). Interestingly, novel niche-ALL interactions were identified, such as MIF - (CD74 + CXCR4) for early mesenchymal progenitors and Ephrin A5 - EPHA7 for adipogenic progenitors.

To validate these findings, we performed co-culture studies using patient-derived B-ALL cells with the two stromal populations. Consistent with our computational analysis, we observed that both stromal populations supported *ex vivo* survival of primary ALL cells. However, adipogenic progenitors exhibited a slightly stronger supportive effect compared to early mesenchymal progenitors (Figure 5E). Dose-response analysis of dexamethasone showed that both stromal populations provide chemoresistance to B-ALL cells (Figure 5F). Multiplex cytokine analysis revealed elevated secretion of CXCL12 and osteopontin in the co-culture medium of both stromal populations (Figure 5G & H). Notably, interleukin 7 and stem cell factor were exclusively detected in the co-culture medium of adipogenic progenitors, aligning with our transcriptomic findings (Figure 5I & J).

### Interleukin 7 and VCAM1 are adipogenic progenitor-specific ALL supportive signals

Given the distinct communication pathways identified through transcriptomic analysis by which the two stromal populations support ALL cells, we sought to further investigate these mechanisms *ex vivo*. We first examined whether the enhanced ALL cell survival observed in co-culture with stromal populations was dependent on cytokine signaling. To this end, we targeted the signaling mediated by interleukin-7, osteopontin, and CXCL12 using blocking antibodies against IL7 and osteopontin, and plerixafor, an inhibitor of the CXCL12 receptor CXCR4. While Plerixafor and anti-osteopontin reduced the supportive effect of both stromal populations on ALL cells, anti-IL7 specifically decreased ALL cell survival in co-culture with adipogenic progenitors, without affecting their survival in co-culture with early mesenchymal progenitors or in monoculture (Figure 6A-C). These findings align with the specific secretion of IL7 predominantly by adipogenic progenitors, as shown in Figure 5J. Furthermore, the addition of these cytokines to ALL cell cultures without stromal cell co-culture demonstrated that IL7 alone increased ALL cell survival, whereas osteopontin and CXCL12 did not (Supplementary Figure 10).

**Figure 6.**
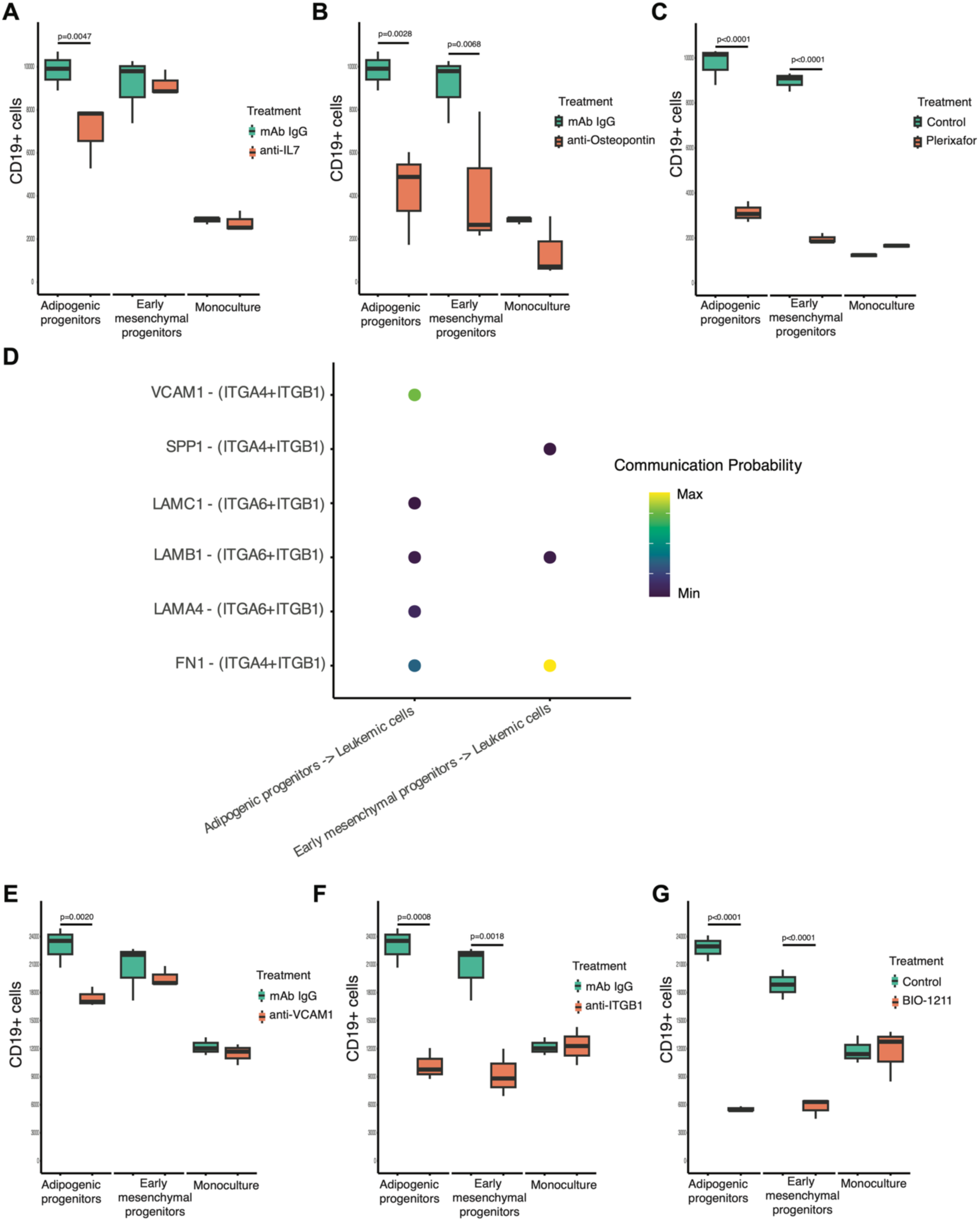
Adipogenic progenitor-specific support of B-ALL cells. Effect of cytokine signaling blockade on ALL cell survival in co-culture with stromal populations. ALL cells were co-cultured with early mesenchymal progenitors, adipogenic progenitors, or cultured alone. Cell survival was assessed after treatment with blocking antibodies against IL7 (A), Osteopontin (B), or the CXCR4 inhibitor Plerixafor (C). D) Ligand-receptor interaction analysis between leukemic cells, adipogenic progenitors and early mesenchymal progenitors. Shown are interactions specifically involving integrin β1 (ITGB1). Dot color reflects interaction probability (from minimum to maximum). E) Effect of VCAM1 blockade on ALL cell viability in co-culture or monoculture. F-G) Effect of blocking integrin beta 1 (ITGB1) using a blocking antibody (F) or the small peptidomimetic drug BIO-1211 (G) on ALL cell viability in co-culture or monoculture.

Our cell-cell communication analysis revealed that the VCAM1/(ITGA4-ITGB1) axis was exclusively utilized by adipogenic progenitors to interact with ALL cells (Figure 6D). Blocking this interaction in *ex vivo* culture with a VCAM1 blocking antibody led to a decreased viability of leukemic cells in co-culture with adipogenic progenitors, but not with early mesenchymal progenitors or in monoculture (Figure 6E). In contrast, blocking antibodies against integrin beta 1 (ITGB1) or the small peptidomimetic drug BIO-1211, which targets the integrin beta 1 binding site, effectively decreased the viability of leukemic cells in co-culture with both stromal populations (Figure 6F & G). This broader effect of ITGB1 blockade is consistent with its involvement in other cell-cell interaction axes with stromal populations, including extracellular matrix proteins as partners, as observed in Figure 6D.

### B-ALL-induced adipogenic bias in stromal cells

Differential abundance analysis revealed a significant increase in the proportion of adipogenic progenitors in B-ALL patient-derived stromal cells compared to healthy donor-derived cells (Figure 7A & B), suggesting a bias towards the adipogenic lineage in patients with B-ALL. To investigate whether this bias is influenced by the leukemic environment, we assessed the adipogenic differentiation potential of unsorted ex vivo expanded stromal cells derived from B-ALL patients and healthy donors. Our results demonstrated that B-ALL-derived stromal cells exhibited a significantly higher capacity for adipogenic differentiation compared to healthy donor-derived cells (Figure 7C & D). To further explore the underlying mechanisms, we co-cultured healthy donor-derived mesenchymal stromal cells with patient-derived B-ALL cells. We observed that exposure to leukemic cells enhanced MSC differentiation toward the adipogenic lineage (Figure 7E & F), indicating that leukemia-specific factors can directly reprogram stromal cells. Notably, this adipogenic skewing was dependent on cell-cell contact, as evidenced by the lack of enhanced adipogenic differentiation when B-ALL cells and MSCs were cultured in a transwell system (Figure 7G). Collectively, these findings indicate that B-ALL can induce a contact-dependent shift in stromal cell differentiation towards the adipogenic lineage.

**Figure 7.**
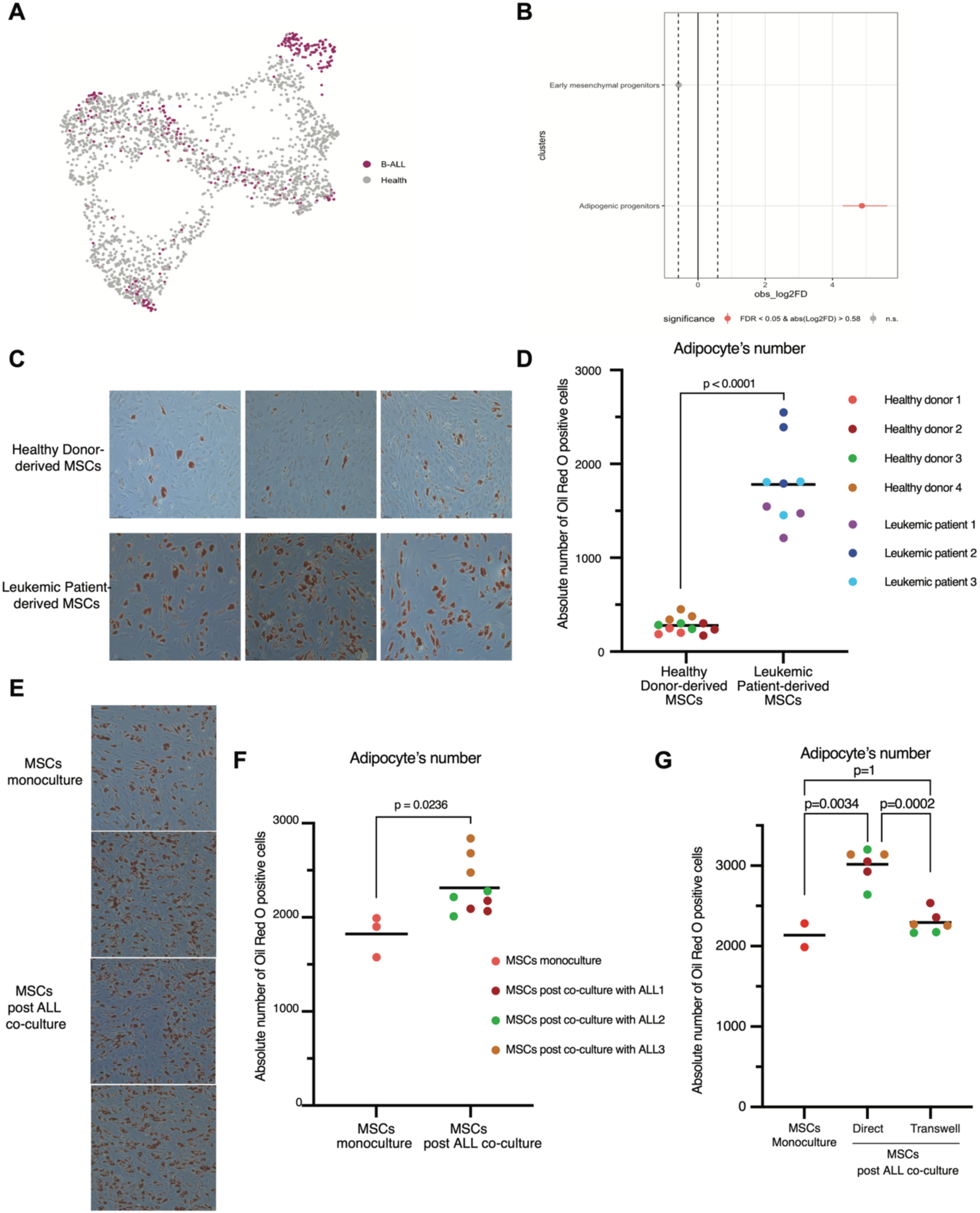
A) UMAP representation of stromal cell clusters from healthy (grey) and ALL patients (violet). B) Point-range plot showing the pairwise proportional difference for each stromal cell cluster (Monte Carlo/permutation test). Red dots indicate statistically significant differences (FDR < 0.05), and vertical dashed lines mark the absolute value of the log2 fold difference (FD) cutoff for significance. C) Representative images of stromal cells derived from three healthy donors (first row) and three leukemic donors (second row) cultured in adipogenic medium for 14 days. Oil red O stains lipid droplets (red). D) Quantification of lipid droplet-positive cells per well in a 96-well plate. Data are presented as mean ± standard deviation. Statistical significance was assessed by a two-tailed unpaired Student’s t-test. E) Representative images of stromal cells post-monoculture (first column) and post-co-culture with three patient-derived xenografts (second, third, and fourth columns) cultured in adipogenic medium for 14 days. Oil red O stains lipid droplets (red). F) Quantification of lipid droplet-positive cells per well in a 96-well plate. Data are presented as mean ± standard deviation. Statistical significance was assessed by a two-tailed unpaired Student’s t-test. G) Average number of droplet-positive cells differentiated from MSCs following co-culture with leukemic cells under direct or trans-well conditions (n=2 wells per condition). Data are presented as mean ± standard deviation. Colors depict different patients, as represented in F. Statistical analysis was performed using a linear mixed-effects model with Condition as a fixed effect and Patient as a random effect to account for inter-patient variability. A significant main effect of condition was found (p = 0.00057). Post-hoc pairwise comparisons with Bonferroni correction are shown in the figure.

### Stromal niche composition and clinical implications

The composition of the stromal niche varied between B-ALL patients (Figure 8A). To assess the prevalence of stromal subpopulations in a larger cohort, we performed gene expression deconvolution analysis using CIBERSORTx on a cohort of bone marrow bulk RNA-seq data from 1374 B-ALL patients (Figure 8B), encompassing a range of different molecular subtypes representing the B-ALL genomic landscape (Figure 8C). The abundance of adipogenic progenitors in bone marrow aspirates did not correlate with leukemic burden (Figure 8D). When stratified by sex, no significant differences were observed. A weak but statistically significant negative correlation with age was detected (R= -0.078, p=0.0058), suggesting a slight decrease in abundance with increasing aging (Supplementary Figure 11). Adipogenic progenitor levels varied across molecular subtypes (Figure 8F). Specifically, the ETV6-RUNX1 subtype exhibited significantly higher levels of adipogenic progenitors compared to BCR-ABL (adjusted p = 0.034), BCR-ABL-like (adjusted p = 1.68e-06), CRLF2r (adjusted p = 6.25e-04), Hyperdiploid (adjusted p = 0.003), iAMP21 (adjusted p = 7.80e-05), and PAX5 (adjusted p = 4.93e-04) subtypes. However, despite statistical significance, the visual differences in abundance across subtypes appear modest, suggesting that factors beyond molecular subtype may more strongly influence stromal microenvironment composition. No significant differences in adipogenic progenitor abundance were observed between risk stratification groups (Figure 8E). Importantly, adipogenic progenitors were significantly enriched in patients experiencing relapse, even after correcting for age, sex and molecular subtype, suggesting their involvement in disease progression (Figure 8G).

**Figure 8.**
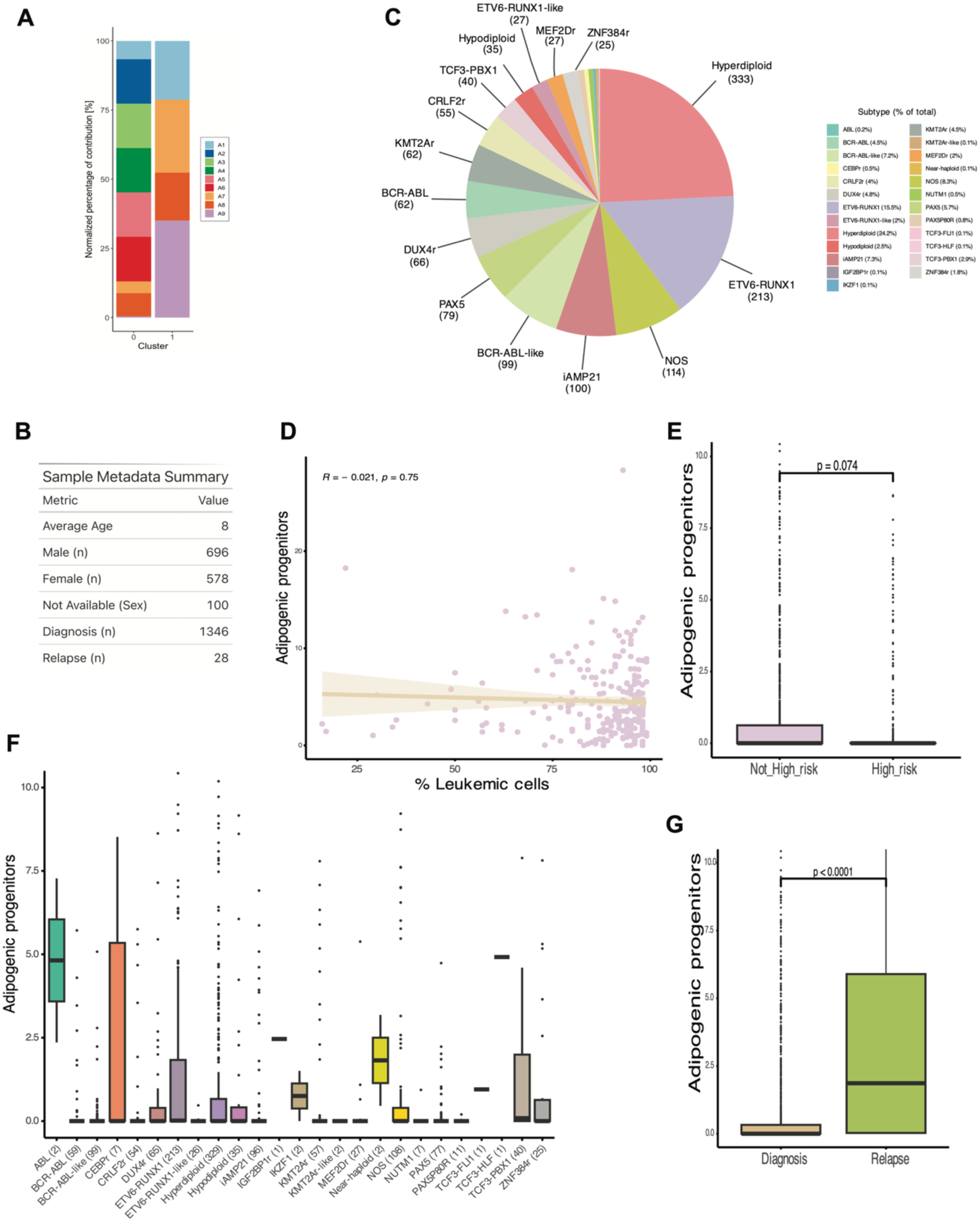
Distinct distribution of stromal cells in B-ALL bone marrow niche. A) Bar plot representing the distribution of stromal cell populations between the different patients analyzed in this study by single-cell RNA sequencing. Early mesenchymal progenitors (cluster 0) and adipogenic progenitors (cluster 1). Each patient sample is represented with a color. B) Sample characteristics of the bulk RNA-seq cohort used for the deconvolution analysis C) Molecular subtype distribution in the cohort. D) Correlation analysis between adipogenic progenitor signature score and the percentage of leukemic cells in bone marrow samples (spearman correlation). E) Comparison of adipogenic progenitor abundance between risk stratification groups. Estimated marginal means (EMMs) were calculated using a linear model adjusting for age and sex (formula: Adipogenic progenitors ∼ Risk Stratification + Age + Sex). Pairwise comparisons between risk groups were performed using the emmeans package, and p-values were derived from the contrast of EMMs. F) Analysis of adipogenic progenitors between different molecular subtypes. A Kruskal–Wallis test was used to assess global differences across subtypes (p = 4.6 × 10⁻¹²). G) Analysis of the adipogenic progenitors in patients at diagnosis and relapse. Estimated marginal means (EMMs) were derived from a linear model adjusting for age, sex, and molecular subtype (Adipogenic progenitors ∼ sample_type + Age + Sex + molecular_subtype). Pairwise comparison between sample types was performed using the emmeans package, and the associated p-value was obtained from the contrast of EMMs.

These findings highlight the heterogeneity of the stromal niche in B-ALL and suggest that adipogenic progenitors may play a crucial role in the leukemic niche.

## Discussion

The bone marrow microenvironment is complex and dynamic and plays a critical role in hematopoiesis and leukemogenesis. Our study provides a comprehensive single-cell transcriptional landscape of the B-ALL niche, revealing a previously unappreciated heterogeneity within the stromal compartment. By identifying distinct early mesenchymal and adipogenic progenitor populations, we have unveiled a more nuanced understanding of the cellular composition of the leukemia-supportive niche.

Our findings challenge the traditional view of a homogenous stromal population in B-ALL and highlight the importance of considering cellular heterogeneity in the stromal niche. The functional characterization of early mesenchymal and adipogenic progenitors revealed distinct properties, with early mesenchymal progenitors exhibiting a more stem-like phenotype and adipogenic progenitors demonstrating enhanced differentiation capacity. These results suggest that the bone marrow stroma is composed of a continuum of cell states, with early mesenchymal progenitors representing a more primitive population that gives rise to adipogenic and potentially other stromal cell types. These findings are consistent with previous studies describing the heterogeneity of bone marrow MSCs in normal hematopoiesis [14, 16, 17, 31, 32]. However, our study provides new insights into the specific functional roles of these subpopulations in the context of B-ALL.

The observed difference in proliferation rate between early mesenchymal and adipogenic progenitors suggests that previous studies have overlooked the heterogeneity of the niche due to the selection bias associated with *ex vivo* propagation of MSCs. The more differentiated adipogenic progenitors are less efficient at expanding in culture, leading to an overrepresentation of early mesenchymal progenitors in *ex vivo*-derived stromal populations. This could have contributed to the oversight of the role of adipogenic progenitors in previous studies.

Additionally, our study reveals that B-ALL induces a significant shift in stromal cell differentiation towards the adipogenic lineage. We observed an increased proportion of adipogenic progenitors in B-ALL patient-derived stromal cells compared to healthy donor-derived cells, suggesting a leukemia-induced adipogenic bias. This bias was further supported by our finding that B-ALL-derived stromal cells exhibited a higher capacity for adipogenic differentiation. Notably, co-culture experiments demonstrated that ALL-derived factors directly reprogram healthy donor-derived stromal cells, enhancing their differentiation toward the adipogenic lineage, in a contact-dependent manner. These results suggest that leukemic cells actively modulate the bone marrow microenvironment, promoting the generation of adipogenic progenitors as part of their supportive niche, which goes in line with previously published studies demonstrating adipocyte-induced skewing of ALL stroma [33]. This adipogenic bias could have profound implications for leukemic progression and therapy resistance, as adipogenic progenitors produce key growth factors and adhesion molecules, such as IL7, SCF, and VCAM1, that support leukemic cell survival.

Our study reveals that there are distinct stromal niches with which ALL cells can interact. More importantly, our data suggest that ALL cells utilize different modes of interaction with these diverse niche cells. We have previously demonstrated the involvement of N-cadherin in leukemia cell support [8], and now confirm that it is expressed on the early mesenchymal progenitors. In contrast, CXCL12, known to support ALL [30, 34], is primarily expressed by adipogenic progenitors. This complexity underscores the challenges in targeting the niche, as leukemic cells can engage with various environments to survive.

A key finding of our study is the identification of adipogenic progenitors as a major component of the leukemic niche. These cells resemble previously described marrow adipogenic lineage precursors (MALPs) and Adipo-CXCL12-abundant-reticular cells (Adipo-CAR) in mice [16, 17]. Our study provides direct evidence for the involvement of adipogenic progenitor cells in supporting leukemic cell growth and their enrichment at relapse. This observation is in line with previous reports highlighting the role of adipocytes in tumor progression [35]. Adipogenic progenitors exhibited high expression of growth factors, such as IL7 and SCF, and adhesion molecules, such as VCAM1, known to support leukemic cell survival, proliferation and therapy resistance [26, 28, 29]. Interestingly, deletion of IL7 in adipogenic progenitors using the Adipoq-Cre transgene mice resulted in severe impairment of B lymphopoiesis [36], suggesting that B cell development relies on adipogenic progenitors. Additionally, deletion of SCF in adipogenic progenitors diminished HSPCs and myeloid progenitors in the bone marrow and led to the development of macrocytic anemia [37], highlighting the crucial involvement of adipogenic progenitors in steady-state hematopoiesis. These findings highlight the broader role of adipogenic progenitors beyond B-ALL and warrant further investigation in other hematological malignancies, such as acute myeloid leukemia.

The heterogeneity of the stromal niche observed in our patient cohort underscores the complexity of the bone marrow microenvironment in B-ALL and highlights the need for a comprehensive understanding of stromal-leukemic interactions to inform the development of effective niche-targeted therapies. Stratifying patient subgroups based on stromal composition may yield additional prognostic insights and help guide future therapeutic approaches. Clearly, studies involving in-depth characterization and spatial mapping of the leukemic microenvironment across larger patient cohorts are essential to fully elucidate the role of distinct niche components in leukemic progression and resistance to both chemo- and immunotherapy in ALL.

In conclusion, our study provides novel insights into the cellular complexity of the B-ALL microenvironment and identifies adipogenic progenitors as a key novel component of the B-ALL niche.

### Limitations of the study

While our study provides novel insights into the B-ALL microenvironment, it is important to acknowledge certain limitations. First, the focus on pediatric patients may limit the generalizability of our findings to adult B-ALL. Second, the sample size of our single-cell RNA sequencing study may not fully capture the variability within the B-ALL patient population. Third, we annotated our pediatric single-cell RNAseq dataset using an atlas derived from total hip arthroplasty samples of adult patients. This might introduce potential caveats due to differences in age, sample source (aspirate versus total tissue) and disease context. Finally, bone marrow biopsies are not routinely performed in children with ALL and, therefore, these analyses rely mostly on bone marrow aspirates. Stroma cell populations that due to their firm attachment to the bone structure are underrepresented in these aspirates, for instance, osteogenic-lineage cells, may add further complexity to the leukemic microenvironment.

## Author contributions

**M.N.F.B.** conceptualized the study, conducted bioinformatic analyses, performed ex vivo experiments, interpreted the data, and drafted and prepared the manuscript. **B.K.** conducted the ex vivo assays presented in Figures 3, 4, and 5, and contributed to manuscript writing and preparation. **M.B.** provided critical feedback on the sorting strategy and assisted in manuscript preparation. **O.H.** contributed to supervision and manuscript preparation. **H.J.V.** oversaw the design, interpretation, and analysis of all experiments and contributed to manuscript writing and preparation.

## Materials and methods

### Bone marrow samples

Bone marrow aspirates were obtained from pediatric B-ALL patients in conjunction with routine clinical sampling after obtaining informed consent from parents and/or legal guardians in accordance with institutional review board approval at the Princess Máxima Center (application number PMCLAB2022.305). Bone marrow mononuclear cells (BM-MNCs) were isolated from fresh bone marrow aspirates using density gradient centrifugation and cryopreserved in liquid nitrogen. Healthy donor-derived bone marrow mononuclear cells used in the single-cell RNA-seq analysis were purchased from Stem Cell Technologies (Catalogue 70001). Patient characteristics, including age, gender, genetic risk factors, and disease stage, can be found in Supplementary Table 1.

### FACs enrichment of stromal, hematopoietic and leukemic cells for scRNAseq

Cryopreserved bone marrow mononuclear cells were thawed and resuspended in RPMI 1640 medium containing 5% FBS and 1 mM EDTA. The cell suspension was passed through a 70 μm mesh and stained for 20 minutes at 4 degrees with an antibody cocktail containing CD19-PE (Catalogue 302207, Clone HIB19, 1:100), CD235a-FITC (Catalogue 349103, Clone HI264, 1:100), CD45-PE-Cy7 (Catalogue 304015, Clone HI30, 1:100), all from Biolegend. Cells were washed twice with medium followed by 7AAD staining (Catalogue 420404, 1:100) to exclude dead cells before sorting on a Sony SH800S instrument (Sony Biotechnology). Leukemic cells, hematopoietic, and non-hematopoietic cells were sorted in RPMI 1640 medium containing 5% FBS. Samples were kept during the whole procedure at 4 °C. Following enrichment, each of the three cell fractions (leukemic, hematopoietic, non-hematopoietic) were resuspended at a concentration of 1000 cells/μL in PBS + 0.04% BSA. Cells were pooled in a 1:1:1 ratio and the cell suspension was subsequently mixed with a cell suspension from another patient with a different gender (multiplexed boy/girl samples). Subsequently, the cell suspension was loaded into the 10x Genomics Chip as described in ‘Single-cell RNA library generation’.

### Single-cell RNA-seq library generation

In total, 20,000 cells were loaded for each (multiplexed) sample onto a 10x Genomics Chip G. We used Chromium Next GEM Single Cell 3’ GEM, Library & Gel Bead Kit and followed the protocol as provided by 10x Genomics. Sequencing was performed on Illumina NovaSeq 6000 sequencer and parameters were set according to 10x Genomics recommendations. A healthy donor sample was included in the scRNA-seq analysis to provide a baseline for comparison. Only the non-hematopoietic compartment from this sample was sorted and sequenced.

### Data processing and filtering

Reads from the sequencing were aligned to the human reference genome (GRCh38) using Cell Ranger v7.1.0. Samples were demultiplexed using souporcell based on Single Nucleotide Polymorphisms (SNPs) identified within the reads. Patient identities were assigned based on specific sex chromosome genes. Further analysis was performed using R v4.0.2 (https://www.r-project.org/) including several packages which are mentioned in the according section. As an initial step, we filtered out low-quality cells. Cells with less than 200 expressed features were removed. Additionally, cells with greater than 10% of reads mapping to mitochondrial genes were excluded. Doublets/multiplets, which can arise during library preparation, were identified and removed using the scDblFinder package with default parameters [38].

### Data integration

All patient scRNAseq datasets were merged into one Seurat object after prefiltering (see section: “Data processing and filtering”). Based on this object, the data was normalized using the SCTransform function from Seurat with default parameters. Principal component analysis (PCA) was performed using the RunPCA function from Seurat, with the number of principal components (npcs) set to 30, a value determined empirically to capture the major sources of variation in the data. The IntegrateLayers function was run with the RPCA integration method and default settings to integrate the data across different patients and account for batch effects.

### Annotation

Cluster annotation for the full integration of all samples was performed at a low-resolution to identify major cell types. Cell types were annotated based on information from literature, functional information from gene sets based on the marker genes of a cluster and unbiased cell type recognition using the Human Primary Cell Atlas and bone marrow scRNAseq references [14, 15] in SingleR with default parameters [39]. The cell type annotation of the stromal cluster relied primarily on the dataset by Bandyopadhyay et al. as a reference in SingleR using default parameters. Unsupervised clustering of the stromal cluster using the FindClusters function in Seurat identified 5 subclusters. However, the biological relevance and distinct cell identities of these subclusters could not be elucidated based on their top markers. Furthermore, the transcriptome of 4 of these subclusters exhibited greater similarity to each other compared to the fifth cluster. To further investigate the stromal cell populations, single-cell RNA-seq data from the manuscript of Bandyopadhyay et al, Cell 2024 [14], was downloaded from NCBI Gene Expression Omnibus (GSE253355) and loaded into Seurat. This comparative analysis predominantly identified two stromal populations: early mesenchymal progenitors (fibro-MSCs) and adipogenic progenitors (adipo-MSCs).

### Marker gene identification

Marker genes per cluster were calculated by the Seurat function FindAllMarkers, using the method “Model-based Analysis of Single-cell Transcriptomics” (MAST) with a log fold change threshold of 0.3 and min-pct set to 0.3. Enriched gene sets per set of marker genes were analyzed as described in the Gene Set Enrichment Analysis section.

### Gene set enrichment analysis (GSEA)

The gprofiler2 package was used to perform GSEA based on marker genes and differentially expressed genes [40]. The gost function was run with default settings. Ribosomal genes were excluded from the gene lists beforehand to focus on biologically relevant processes and only gene lists with more than 5 remaining genes sorted by adjusted p value were tested. Before plotting, the identified gene sets were filtered for terms with less than 1000 genes to exclude overly broad terms. Only terms from GO, Reactome and KEGG were plotted.

### Signal pathway activity

To investigate signaling pathway activity across stromal cell populations, we utilized the R package PROGENy [41]. The integrated Seurat object was used to annotate stromal subpopulations and compute pathway activity scores using the top 500 genes relevant for human signaling pathways. PROGENy scores were added to the Seurat object as a new assay and scaled using Seurat’s ScaleData function. Pathway scores were extracted and aggregated by cell type to compute average activity levels per population. Visualization of global and selected pathway activities was performed using heatmaps (pheatmap package), displaying the average pathway activity per cell type.

### Transcription factor activity inference using SCENIC

To infer transcription factor (TF) activity across stromal populations in B-ALL, we applied the SCENIC (Single-Cell rEgulatory Network Inference and Clustering) pipeline [18].. Raw gene expression counts were extracted from an integrated Seurat object containing stromal cells. SCENIC was executed in R using the GENIE3, RcisTarget, and AUCell packages with human motif annotations and cisTarget databases (hg19) for regulatory network inference. Regulon activity was summarized per stromal cluster and visualized using heatmaps to identify cell type-specific TF programs.

### Differentiation state prediction

To infer the differentiation potential of stromal cells, we employed CytoTRACE v0.3.3, a computational tool that estimates developmental states based on gene expression profiles and transcriptional entropy [42]. We began by extracting the raw count matrix from the RNA assay of the Seurat object. This count matrix was converted into a data frame suitable for input into the CytoTRACE algorithm. CytoTRACE was then run using default parameters, returning a differentiation score for each cell, where higher scores reflect a more progenitor-like state and lower scores indicate greater differentiation. The CytoTRACE scores were subsequently added to the Seurat object as metadata. Ridge plots were created to characterize the density of differentiation states across cell types.

### Cell-cell communication analysis

The CellChat algorithm was applied to perform an unbiased ligand-receptor interaction analysis [43]. CellChat’s human ligand-receptor database was employed to infer potential cell-cell interactions. Overexpressed genes and interactions were identified, followed by projection onto a human protein-protein interaction network. Communication probabilities were computed using a tri-mean method, and signaling pathways were aggregated and filtered to include interactions supported by at least 10 cells per group. Network centrality metrics were calculated, and interaction patterns were visualized via circle plots and bubble plots (targeted to key populations such as adipogenic and early mesenchymal progenitors, and leukemic cells).

A leukemic cell interaction score was calculated per cell cluster as the average expression of signaling factors (cytokines, integrins, extracellular membrane components) that were inferred to interact with leukemic cells based on the CellChat inferred interactions, using a curated gene set of 45 nourishing ligands (e.g., CXCL12, FN1, VEGFA, GDF15). Module scores were calculated per cell, and statistical differences across non-leukemic cell types were evaluated using Kruskal-Wallis and post-hoc Bonferroni-adjusted Wilcoxon tests.

To identify dominant modes of outgoing intercellular communication, non-negative matrix factorization (NMF) was applied to the signaling networks. The optimal number of outgoing signaling patterns was determined using the selectK() function, which evaluates cophenetic and silhouette coefficients; both metrics indicated an elbow at k = 5, suggesting five distinct communication programs. These five communication patterns represent core signaling strategies employed by different cell types across the microenvironment. Each pattern groups cell populations that exhibit similar outgoing signaling behaviors, highlighting functional convergence or specialization.

### Inference of copy number variation

To distinguish leukemic cells from non-malignant populations in pediatric B-ALL samples, we performed copy number alteration (CNA) analysis on individual patients using the SCEVAN (Single-Cell Evolutionary Variational Autoencoder for CNAs) R package [44]. For each patient, we extracted single-cell gene expression data from an integrated Seurat object. To define a set of reference "normal" cells for CNA inference, clusters corresponding to T cells and monocytes were identified based on expression of canonical markers such as CD3G, CD8A, CD14, and FCGR3A, using DotPlot() visualization. Cells from clusters expressing these markers were selected as the normal reference population. Gene expression matrices were extracted from the RNA assay for each sample and converted to matrix format for input into the pipelineCNA() function from the SCEVAN package. SCEVAN output classified cells based on inferred CNA profiles into likely malignant or normal categories.

### Spatial transcriptomic analysis

We analyzed a publicly available spatial transcriptomics dataset generated using the Xenium In Situ platform (10x Genomics) from a formalin-fixed paraffin-embedded (FFPE) trephine bone marrow biopsy sample obtained from a patient diagnosed with B-cell acute lymphoblastic leukemia (B-ALL) [25]. Patient metadata, including age, sex, and molecular subtype, were not provided with the dataset. The dataset consisted of a total of 225,906 spatially resolved cells. The median number of transcripts per cell was 27, and a total of 6,933,297 high-quality decoded transcripts were detected across the tissue section. The total imaged region area was 26,288,545.2 µm². Gene expression profiling was performed using a panel of 477 RNA targets, which included 377 predesigned genes from the standard Xenium Human Multi-Tissue Panel and 100 custom-selected genes relevant to bone marrow.

Cell segmentation was performed using the Xenium onboard pipeline. Nuclear boundaries were identified based on DAPI staining, and a heuristic expansion was applied to approximate full cell boundaries, enabling transcript assignment to individual cells. Xenium data was analyzed using the Seurat (v5) R packages. Xenium raw data were imported with LoadXenium() and filtered to retain cells with >10 UMIs and >3 detected genes. Quality control metrics were visualized using violin plots. Selected regions of interest were cropped for high-resolution spatial inspection using the Crop() function. Normalization was performed using SCTransform. Principal component analysis (PCA), uniform manifold approximation and projection (UMAP), and shared nearest neighbor-based clustering were used for dimensionality reduction and cell clustering (RunPCA(), RunUMAP(), FindNeighbors(), FindClusters()), with clustering resolution set to 0.2 for the full dataset. Marker genes for each cluster were identified via differential expression analysis using the MAST test (FindAllMarkers()), applying a threshold of log2FC > 0.1 and adjusted p-value < 0.05. To identify and map specialized cellular microenvironments, spatial niches were inferred using the BuildNiche()function from Seurat with standard settings, which detects spatially enriched co-localized cell populations. For spatial visualization, annotated cell identities were exported and visualized using Xenium Explorer 3.0.

### Generation of single-cell reference matrix

A single-cell RNA-seq reference was constructed using our integrated dataset encompassing all cellular populations within the B-ALL niche. A reference expression matrix was generated by extracting raw count data from the RNA assay using GetAssayData() with the "counts" layer. The resulting matrix was converted to a data.frame and matched with corresponding cell-type annotations. The column names of the expression matrix were relabeled with their respective "Annotation_L2" identities to ensure compatibility with CIBERSORT, which requires labeled expression profiles as input. The final reference matrix was saved as a tab-delimited text file (.txt) for use in downstream deconvolution of bulk RNA-seq data.

### Cell type deconvolution Analysis in Bulk RNA-seq Data

Bulk RNA-seq data generated for routine diagnostics were obtained from the Princess Máxima Center Biobank (application number PMCLAB2021.258), containing a cohort of 245 B-ALL bone marrow patient samples. Furthermore, we obtain a publicly available dataset downloaded from the St. Jude Cloud Platform via DNAnexus [45]. For RNA-seq deconvolution, we employed CIBERSORTx with the Impute Cell Fractions job type, with batch correction S-mode, in absolute mode and 100 permutations. Clinical metadata including age, sex, molecular subtype, and sample type (diagnosis vs. relapse) were imported into R for further analysis. Samples were grouped into risk categories (High vs. Not High Risk) based on the molecular subtype according to criteria described in the literature [46], and annotated accordingly. Samples were categorized as High Risk if they harbored one of the following subtypes: BCR-ABL, BCR-ABL-like, KMT2A rearranged (KMT2Ar), hypodiploid, near-haploid, MEF2D rearranged (MEF2Dr), MYC, iAMP21, TCF3-HLF, or ABL-class fusions. Samples were categorized as Not High Risk if they exhibited one of the following subtypes: ETV6-RUNX1, ETV6-RUNX1-like, hyperdiploid, TCF3-PBX1, DUX4 rearranged (DUX4r), CRLF2 rearranged (CRLF2r), CEBP rearranged (CEBPr), KMT2A-like, ZNF384-like (ZNF384r-like), PAX5, ZNF384 rearranged (ZNF384r), IGF2BP1 rearranged (IGF2BP1r), IKZF1, or other. Outlier detection was performed on the adipogenic progenitor scores using visual inspection of histograms and Grubbs’ test. One extreme outlier (value >300) was identified and excluded from downstream analyses.

### Stromal populations cell sorting

MSC subtypes were isolated from BM-MNCs using fluorescence-activated cell sorting (FACS) on a Sony SH800S instrument (Sony Biotechnology) based on surface marker expression identified from the single-cell RNA sequencing data. In addition to the analysis of well-known stromal CD markers, we performed an unbiased approach using the sc2marker package with default settings [47].

To exclude hematopoietic and leukemic cells, we used a dump gate strategy at 421 nm emission composed of CD19-BV421 (Catalogue 302233, clone HIB19, 1:100), CD22-BV421 (Catalogue 302523, clone HIB22, 1:100), CD45-BV421 (Catalogue 304031, clone HI30, 1:100), CD235a-BV421 (Catalogue 349131, clone, HI264 1:100). Live/dead exclusion was performed using Zombie green (Catalogue 423111, 1:100). Adipogenic progenitors were sorted as positive for CD106-PE (Catalogue 305805, clone STA, 1:100) and Cadherin 11-APC (Catalogue 368705, clone 16G5, 1:100), while early mesenchymal progenitors were negative for these markers and positive for CD81-PE-Cy7 (Catalogue 349511, clone 5A6, 1:100) and CD90-AF700 (Catalogue 328119, clone 5E10, 1:100). Stromal populations were sorted in MSC medium (DMEM-low glucose supplemented with 10% FBS, 1% Primocin solution and 1 ng/mL bFGF).

### scRNA seq analysis of early mesenchymal progenitors and adipogenic progenitors

Sorted MSC subtypes were cultured in MSC medium (DMEM-low glucose supplemented with 10% FBS, 1% Primocin solution and 1 ng/mL bFGF) and incubated at 37 °C with 5% CO2. Cells were expanded to passage 3 and scRNAseq analysis was performed as described in ‘Single-cell RNA library generation’.

### Projection of sorted stromal populations into the unsorted stromal scRNAseq dataset

The two stromal populations were isolated via FACS from leukemic donor samples and sequenced independently. To control for sampling bias, each sorted population was downsampled to 2,300 cells. Each dataset was normalized using the SCTransform function, regressing out mitochondrial content. Following normalization, the datasets were merged and dimensionality reduction was performed using PCA (30 components), followed by UMAP. The merged dataset of sorted stromal cells was projected onto the UMAP embedding of the reference stromal dataset using the RunKNNMap function from the SCP package. The projection utilized 3,000 integration features identified via SelectIntegrationFeatures, and both query and reference datasets used the SCT assay. Following projection, cells from the sorted populations were classified according to the nearest neighbors in the reference using RunKNNPredict. Cells were assigned to predicted stromal subtypes with a low-frequency filter set at 20 cells per type. To quantify enrichment of predicted stromal subtypes within each sorted population, classification outcomes were tabulated and expressed as proportions. A contingency table comparing the frequencies of adipogenic versus early mesenchymal progenitors across the two sorted populations was constructed. A chi-square test of independence was used to evaluate differences in predicted subtype distributions between sorted populations. Proportional distributions of predicted stromal subtypes were visualized using stacked bar plots in ggplot2, with custom color palettes for clarity.

### Flow cytometry analysis of sorted early mesenchymal progenitors and adipogenic progenitors

Sorted MSC subtypes were cultured in MSC medium (DMEM-low glucose supplemented with 10% FBS, 1% Primocin solution and 1 ng/mL bFGF) and incubated at 37 °C with 5% CO2. Cells were expanded to passage 3, trypsinized and prepared for flow cytometry analysis. The cell suspension was stained with Viakrome 808 (Catalogue #C36628, 1:100, Beckman Coulter) for 15 minutes at 4 degrees to exclude dead cells. After staining, 1 mL of MSC medium was added to the suspension, which was then centrifuged. The supernatant was discarded, and the cell pellet was resuspended in a single marker antibody solution. All antibodies were obtained from Biolegend. Following antibody staining, cells were washed with PBS containing 1% FBS, centrifuged and the supernatant was discarded. The final cell pellet was resuspended in 100 microliters of PBS containing 1% FBS fow flow cytometry analysis on a Cytoflex LX instrument (Beckman Coulter). Data were analyzed using FlowJo software (BD Biosciences).

### Colony-forming unit fibroblast (CFU-F) assays

Sorted cells were seeded in 6-well plates at a density of 475 cells per well (50 cells/cm2) containing culture medium (DMEM-low glucose supplemented with 10% FBS, 1% Penicillin/Streptomycin solution and 1 ng/mL bFGF) and incubated at 37 °C with 5% CO2. Medium was changed every 3–4 days. At day 10, cells were fixed with methanol and stained with 0,5% crystal violet staining solution. Adherent colonies with more than 50 cells were quantified.

### Multipotency assays

Adipogenic and osteogenic differentiation assays were performed in cells at passage three to five. For adipogenic differentiation, cells were seeded at a density of 21,000 cells/ cm2 and cultured in DMEM-low glucose (Gibco, Catalogue 21885-025) supplemented with 10% FBS, 1% Penicillin/Streptomycin, 0.5 μM isobutylmethylxanthine (Sigma, I5879), 60 μM indomethacin (Sigma, 17378), 5 μg/mL insulin (Sigma, I9278) and 1 μM dexamethasone (Sigma, D2915) for 15 days (medium was changed every 3-4 days). To measure adipogenic differentiation, cells were fixed with 4% PFA and stained using 0.5% Oil Red O (Sigma, 00625) in isopropyl alcohol diluted with distilled water in 3:2 ratio and the lipid-filled cells across the whole well of a 96-well plate were counted. Images of stained cells from corresponding samples in 12-well plates were captured for visualization.

For osteogenic differentiation, cells were seeded at a density of 3000 cells/ cm2 in osteogenic medium for 15 days (medium was changed every 3-4 days). The osteogenic differentiation medium contained DMEM supplemented with 10% FBS, 1% Penicillin/Streptomycin, 1% glutamine, 50 μg/mL L-ascorbate acid (Sigma, A8960), 10 mM β-glycerophosphate (Sigma, G6376) and 100 nM dexamethasone. Osteogenic differentiation was verified by alizarin red S staining (Sigma, A5533) by visualization of mineralized deposits.

### Leukemic cell viability in co-culture with stromal populations

Early mesenchymal progenitors and adipogenic progenitors between passages 5-7 were seeded at a cell density of 15,000 cells/cm2 in 96 well plate in MSC medium and incubated at 37 °C with 5% CO2. The next day, 200,000 patient-derived bone marrow mononuclear cells and sorted MSC subtypes were co-cultured in SFEM medium containing 1% Penicillin/Streptomycin solution and incubated at 37 °C with 5% CO2. After 7 days, all cells per well were harvested by aspirating the supernatant and trypsinizing attached cells using TrypLE Express (Gibco, 12605). Both cell suspensions were combined, per well. After washing the cells with PBS, FBS 5%, cells were stained with CD19-BV421 (Biolegend, Catalogue 302233, clone HIB19, 1:100) and Zombie NIR (Biolegend, Catalogue 423106, 1:200) for live/dead exclusion.

Cell viability was assessed using a flow cytometry Cytoflex S (Beckman Coulter), recording all events with a fixed volume of 80 microliters per well.

### Evaluation of small molecules and blocking antibodies in co-culture of B-ALL cells with stromal populations

Early mesenchymal progenitors and adipogenic progenitors between passages 5-7 were seeded at a cell density of 15,000 cells/cm2 in 96 well plate in MSC medium and incubated at 37 °C with 5% CO2. The next day, 200,000 patient-derived bone marrow mononuclear cells were added per well in 200ul SFEM containing 1% Penicillin/Streptomycin. Where indicated, cultures were treated with varying dexamethasone (Sigma, D4902) concentrations (1 nM, 3 nM, 5 nM, 10 nM, 30 nM, 50 nM, 100nM), mouse IgG1 kappa Isotype control (1 ug/mL, MedChemExpress, Catalogue HY-P99977), anti-mouse/human IL-7 (1 ug/mL, BioCell, Clone M25, Catalogue BE0048), Plerixafor (1 nM, MedChemExpress, Catalogue HY-10046), anti-Osteopontin (1 ug/mL, BE0382, BioCell), BIO-1211 (50uM, MedChemExpress, Catalogue HY-14126), anti-hVCAM1 mouse monoclonal IgG (2 ug/mL, R&D Systems, Clone BBIG-V1, Catalogue BBA5), and anti-Integrin Beta1 (2 ug/mL, Merk, Clone AIIB2, Catalogue MABT409). After 7 days of incubation, all cells per well were harvested by aspirating the supernatant and trypsinizing attached cells using TrypLE Express (Gibco, 12605). Both cell suspensions were combined, per well. After washing the cells with PBS, FBS 5%, cells were stained with CD19-PE (Biolegend, Catalogue 302208, Clone HIB19, 1:100) and Zombie Violet Fixable Viablity Kit (BioLegend, Catalogue 423113/423114, 1:100) for live/dead exclusion. The flow cytometry analysis on a Cytoflex LS instrument (Beckman Coulter) was used to measure absolute number of viable CD19+ leukemic cells per well. Data was analyzed with FlowJo software (BD Biosciences).

### Isolation of pediatric bone marrow stromal cells from healthy and leukemic patients

Healthy donor-derived stromal cells were isolated from surplus iliac crest bone chip material harvested from pediatric patients undergoing alveolar bone graft surgery. All human samples were obtained with the approval of the Erasmus MC, University Medical Center Medical Research Ethics Committee (MEC-2014-16). Iliac crest bone chips were washed with expansion medium composed of Minimum Essential Medium (MEM)-α (containing nucleosides) supplemented with heat inactivated 10% v/v fetal bovine serum (FBS) (both Thermo Fisher Scientific, Waltham, MA, USA), 1.5 µg/ml fungizone (Gibco), 50 µg/ml gentamicin (Gibco), 25 µg/ml L-ascorbic acid 2-phosphate (Sigma-Aldrich, St. Louis, MO, USA), and 1 ng/ml fibroblast growth factor-2 (Instruchemie, Delfzijl, The Netherlands), and the resulting cell suspension was seeded in T75 flasks. Cells were washed twice with phosphate buffered saline (Thermo Fisher Scientific) supplemented with 2% v/v heat inactivated FBS 24 hours following seeding to remove non-adherent cells. Stromal cells were cultured at 37°C and 5% carbon dioxide under humidified conditions, with expansion medium refreshed every 3–4 days. A similar approach was used to isolate stromal cells from bone marrow aspirates from B-ALL patients (application number PMCLAB2022.305). 24 hours after seeding, the non-attached cells were removed and adherent cells were cultured at 37°C and 5% carbon dioxide under humidified conditions, with expansion medium refreshed every 3–4 days. Stromal cells were sub-cultured upon reaching 80–90% confluency using 0.25% w/v trypsin-EDTA (Thermo Fisher Scientific) and reseeded at a cell density of 2,300 cells/cm2. Adipogenic differentiation was performed with leukemic and healthy donor-derived stromal cells at passage 5 as described in the section ‘Multipotency assays’.

### Analysis of adipogenic differentiation skewing by leukemic cells

Healthy donor -derived stromal cells were co-cultured with patient-derived xenografts in SFEM medium containing 1% Penicillin/Streptomycin solution and incubated at 37 °C with 5% CO2. After 24 hours, leukemic cells were removed by aspirating the supernatant. After washing the cells with PBS, FBS 5%, adipogenic differentiation was induced and quantified as described in the section ‘Multipotency assays’.

In the same manner, 12-well hanging cell culture inserts, PET membrane, with a pore size of 0.4 μm (cellQART, 9310402) were used for the transwell assay. Three days later, leukemic cells were removed, and the remaining cells were cultured in adipogenic differentiation media for 12 days to match 15 days of adipogenic differentiation period. Five pictures per well were taken at 2X magnification and based on that the average number of lipid containing cells/well was calculated.

### Multiplex cytokine analysis

Harvested supernatant (25 μL) was used for cytokine quantification using a bead-based immunoassay according to the manufacturer’s instructions (LEGENDplex, BioLegend, Catalogue 741362 and 740611). Specific antibody coated beads were incubated with supernatant from each condition, in triplicate, forming an analyte-antibody complex. After washing, a biotinylated detection antibody cocktail was added which bound to the specific analyte-antibody complexes. Streptavidin-phycoerythrin was subsequently added which bound to the biotinylated detection antibodies, providing fluorescent signal intensities in proportion to the bound analyte amount. Fluorescent signals were measured using Cytoflex LX and analysed using LEGENDplex data analysis software where concentrations of each analyte were determined using a standard curve generated from the same assay.

### Statistical Analysis

Unless otherwise stated, statistical significance was assessed by a two-tailed Student’s t-test for comparison between two groups or two-way ANOVA with Bonferroni’s multiple comparison for comparisons among multiple groups. A p value < 0.05 was considered statistically significant. Statistical analyses were performed using GraphPad Prism 9.0.1 and R. GSEA is performed with the hypergeometric test for overrepresentation (one-sided), followed by multiple comparison correction using the Benjamini-Hochberg method. The individual method of correction for specific R packages are described in the corresponding publications.

## Data availability

The single-cell RNA sequencing data generated in this study have been deposited in Zenodo at <u>10.5281/zenodo.13755294</u>. All other relevant data supporting the key findings of this study are available within the article and its Supplementary Information files or from the corresponding author upon request.

## Code availability

Custom scripts are available at: https://github.com/mauriferrao/ALL_Stromal_Microenvironment.git Any additional information required to reanalyze the data reported in this paper is available from the corresponding author upon request.

## Acknowledgements

We would like to thank the single cell sequencing facility, the flow cytometry facility at the Princess Máxima Center, Polina Derevianko, Daniel Mata Casimiro, Anita van Oort and Elizabeth Schweighart for their assistance.

## References

1. Ding, L. and S.J. Morrison, Haematopoietic stem cells and early lymphoid progenitors occupy distinct bone marrow niches. Nature, 2013. 495(7440): p. 231–5.

2. Mendez-Ferrer, S., et al., Mesenchymal and haematopoietic stem cells form a unique bone marrow niche. Nature, 2010. 466(7308): p. 829–34.

3. Bianco, P. and P.G. Robey, Skeletal stem cells. Development, 2015. 142(6): p. 1023–7.

4. Kunisaki, Y., et al., Arteriolar niches maintain haematopoietic stem cell quiescence. Nature, 2013. 502(7473): p. 637–43.

5. Greenbaum, A., et al., CXCL12 in early mesenchymal progenitors is required for haematopoietic stem-cell maintenance. Nature, 2013. 495(7440): p. 227–30.

6. Sugiyama, T., et al., Maintenance of the hematopoietic stem cell pool by CXCL12-CXCR4 chemokine signaling in bone marrow stromal cell niches. Immunity, 2006. 25(6): p. 977–88.

7. Duan, C.W., et al., Leukemia propagating cells rebuild an evolving niche in response to therapy. Cancer Cell, 2014. 25(6): p. 778–93.

8. Pal, D., et al., hiPSC-derived bone marrow milieu identifies a clinically actionable driver of niche-mediated treatment resistance in leukemia. Cell Rep Med, 2022. 3(8): p. 100717.

9. Park, C.S., et al., Stromal-induced epithelial-mesenchymal transition induces targetable drug resistance in acute lymphoblastic leukemia. Cell Rep, 2023. 42(7): p. 112804.

10. Witkowski, M.T., et al., Extensive Remodeling of the Immune Microenvironment in B Cell Acute Lymphoblastic Leukemia. Cancer Cell, 2020. 37(6): p. 867–882 e12.

11. Zyla, J., et al., Evaluation of zero counts to better understand the discrepancies between bulk and single-cell RNA-Seq platforms. Comput Struct Biotechnol J, 2023. 21: p. 4663–4674.

12. Sotillo, E., et al., Convergence of Acquired Mutations and Alternative Splicing of CD19 Enables Resistance to CART-19 Immunotherapy. Cancer Discov, 2015. 5(12): p. 1282–95.

13. Witkowski, M.T., et al., NUDT21 limits CD19 levels through alternative mRNA polyadenylation in B cell acute lymphoblastic leukemia. Nat Immunol, 2022. 23(10): p. 1424–1432.

14. Bandyopadhyay, S., et al., Mapping the cellular biogeography of human bone marrow niches using single-cell transcriptomics and proteomic imaging. Cell, 2024. 187(12): p. 3120–3140 e29.

15. Stuart, T., et al., Comprehensive Integration of Single-Cell Data. Cell, 2019. 177(7): p. 1888–1902 e21.

16. Zhong, L., et al., Single cell transcriptomics identifies a unique adipose lineage cell population that regulates bone marrow environment. Elife, 2020. 9.

17. Tikhonova, A.N., et al., The bone marrow microenvironment at single-cell resolution. Nature, 2019. 569(7755): p. 222–228.

18. Aibar, S., et al., SCENIC: single-cell regulatory network inference and clustering. Nat Methods, 2017. 14(11): p. 1083–1086.

19. Gao, H., et al., Early B cell factor 1 regulates adipocyte morphology and lipolysis in white adipose tissue. Cell Metab, 2014. 19(6): p. 981–92.

20. Ambele, M.A., et al., Adipogenesis: A Complex Interplay of Multiple Molecular Determinants and Pathways. Int J Mol Sci, 2020. 21(12).

21. He, J., et al., Dissecting human embryonic skeletal stem cell ontogeny by single-cell transcriptomic and functional analyses. Cell Res, 2021. 31(7): p. 742–757.

22. Aouadi, M., et al., p38MAP Kinase activity is required for human primary adipocyte differentiation. FEBS Lett, 2007. 581(29): p. 5591–6.

23. Bost, F., et al., The extracellular signal-regulated kinase isoform ERK1 is specifically required for in vitro and in vivo adipogenesis. Diabetes, 2005. 54(2): p. 402–11.

24. Richard, A.J. and J.M. Stephens, The role of JAK-STAT signaling in adipose tissue function. Biochim Biophys Acta, 2014. 1842(3): p. 431–9.

25. Acute Lymphoid Leukemia Bone Marrow, In Situ Gene Expression, Xenium Onboard Analysis by Xenium Analyzer. 2024, 10x Genomics.

26. Mendez-Ferrer, S., et al., Bone marrow niches in haematological malignancies. Nat Rev Cancer, 2020. 20(5): p. 285–298.

27. Hughes, A.M., et al., The Bone Marrow Microenvironment in B-Cell Development and Malignancy. Cancers (Basel), 2022. 14(9).

28. Pinho, S., et al., PDGFRalpha and CD51 mark human nestin+ sphere-forming mesenchymal stem cells capable of hematopoietic progenitor cell expansion. J Exp Med, 2013. 210(7): p. 1351–67.

29. Omatsu, Y., et al., The essential functions of adipo-osteogenic progenitors as the hematopoietic stem and progenitor cell niche. Immunity, 2010. 33(3): p. 387–99.

30. Agarwal, P., et al., Mesenchymal Niche-Specific Expression of Cxcl12 Controls Quiescence of Treatment-Resistant Leukemia Stem Cells. Cell Stem Cell, 2019. 24(5): p. 769–784 e6.

31. Baccin, C., et al., Combined single-cell and spatial transcriptomics reveal the molecular, cellular and spatial bone marrow niche organization. Nat Cell Biol, 2020. 22(1): p. 38–48.

32. Wolock, S.L., et al., Mapping Distinct Bone Marrow Niche Populations and Their Differentiation Paths. Cell Rep, 2019. 28(2): p. 302–311 e5.

33. Heydt, Q., et al., Adipocytes disrupt the translational programme of acute lymphoblastic leukaemia to favour tumour survival and persistence. Nat Commun, 2021. 12(1): p. 5507.

34. Spiegel, A., et al., Unique SDF-1-induced activation of human precursor-B ALL cells as a result of altered CXCR4 expression and signaling. Blood, 2004. 103(8): p. 2900–7.

35. Shafat, M.S., et al., Leukemic blasts program bone marrow adipocytes to generate a protumoral microenvironment. Blood, 2017. 129(10): p. 1320–1332.

36. Mukohira, H., et al., Mesenchymal stromal cells in bone marrow express adiponectin and are efficiently targeted by an adiponectin promoter-driven Cre transgene. Int Immunol, 2019. 31(11): p. 729–742.

37. Zhang, Z., et al., Bone marrow adipose tissue-derived stem cell factor mediates metabolic regulation of hematopoiesis. Haematologica, 2019. 104(9): p. 1731–1743.

38. Germain, P.L., et al., Doublet identification in single-cell sequencing data using scDblFinder. F1000Res, 2021. 10: p. 979.

39. Aran, D., et al., Reference-based analysis of lung single-cell sequencing reveals a transitional profibrotic macrophage. Nat Immunol, 2019. 20(2): p. 163–172.

40. Kolberg, L., et al., gprofiler2 -- an R package for gene list functional enrichment analysis and namespace conversion toolset g:Profiler. F1000Res, 2020. 9.

41. Schubert, M., et al., Perturbation-response genes reveal signaling footprints in cancer gene expression. Nat Commun, 2018. 9(1): p. 20.

42. Gulati, G.S., et al., Single-cell transcriptional diversity is a hallmark of developmental potential. Science, 2020. 367(6476): p. 405–411.

43. Jin, S., et al., Inference and analysis of cell-cell communication using CellChat. Nat Commun, 2021. 12(1): p. 1088.

44. De Falco, A., et al., A variational algorithm to detect the clonal copy number substructure of tumors from scRNA-seq data. Nat Commun, 2023. 14(1): p. 1074.

45. Brady, S.W., et al., The genomic landscape of pediatric acute lymphoblastic leukemia. Nat Genet, 2022. 54(9): p. 1376–1389.

46. Malard, F. and M. Mohty, Acute lymphoblastic leukaemia. Lancet, 2020. 395(10230): p. 1146–1162.

47. Li, R., et al., Detection of cell markers from single cell RNA-seq with sc2marker. BMC Bioinformatics, 2022. 23(1): p. 276.

