## Supplementary_Figures for "Distinct Stromal Cell Populations Define the B-cell Acute Lymphoblastic Leukemia Microenvironment"

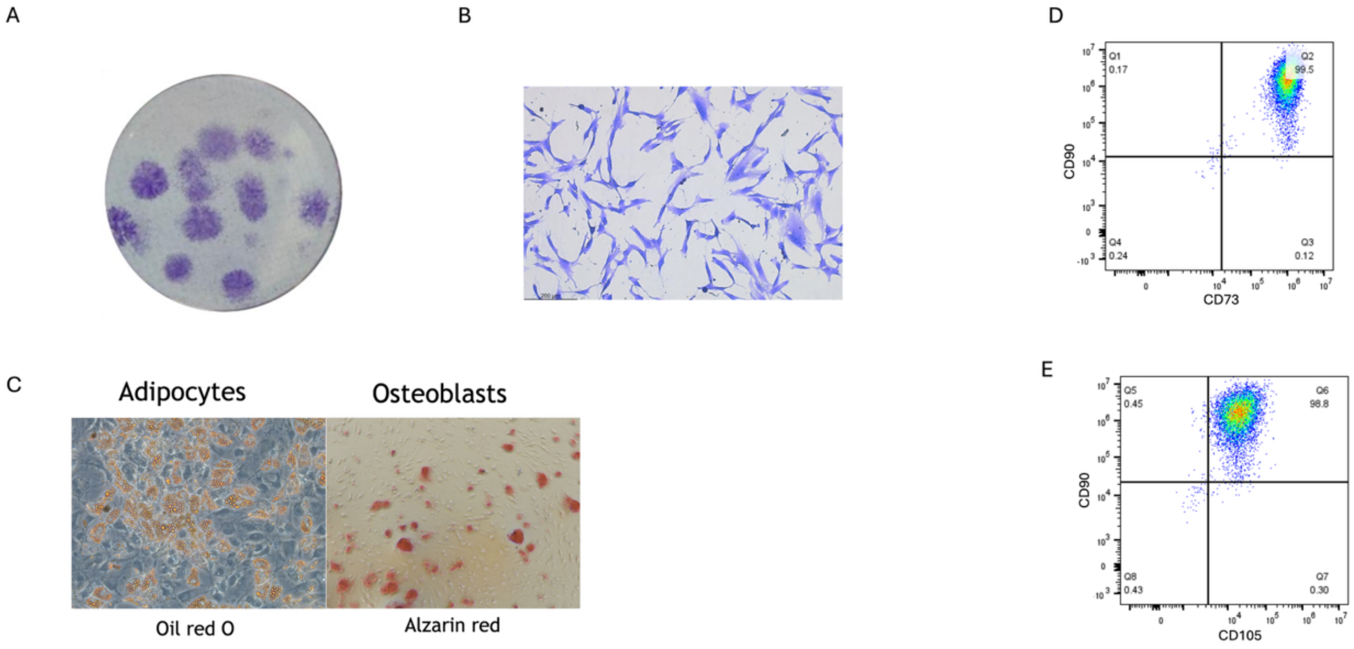


Supplementary Figure 1.

Supplementary Figure 1. Characterization of FACs-sorted viable CD19-CD45-CD235a- bone marrow mononuclear cells after expansion for three passages *in vitro*. A) Colony-forming unit fibroblast (CFU-F) assay stained with crystal violet. A total of 475 cells per well were plated in a 6-well plate (50 cells/cm2). B) Magnification of colony from A, observing fibroblastic-like plastic adherent cells. C) Adipogenic and osteogenic differentiation assay. Oil red O and Alzarin red are used to stain oil droplets and calcium deposition, respectively. D, E) Flow cytometry analysis of CD90, CD73 and CD105 on viable cells.

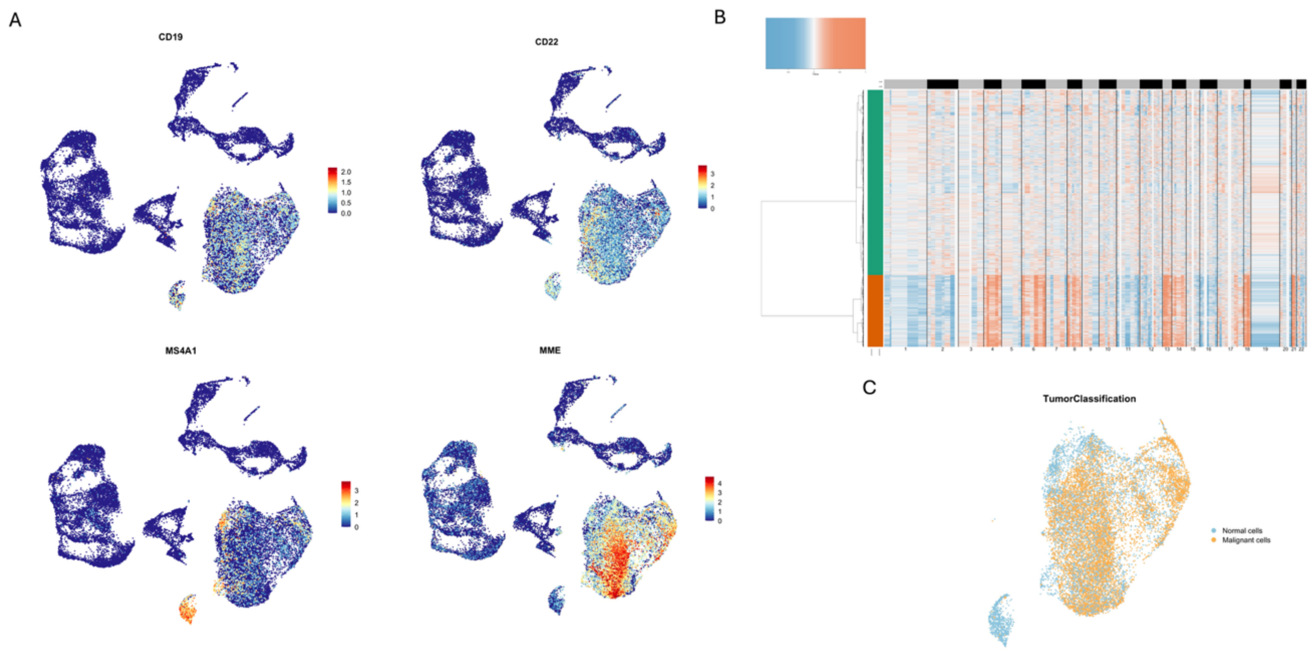


Supplementary Figure 2.

Supplementary Figure 2. Identification of leukemic and healthy B cells A) UMAP representation of B-lineage cell markers. B) Inference of copy number variation (CNV). Heatmap indicates amplifications (red) or deletions (blue) in chromosome regions. C) UMAP representation of the B cell and leukemic cell cluster with the inferred classification as tumor or healthy cells based on CNV’s.

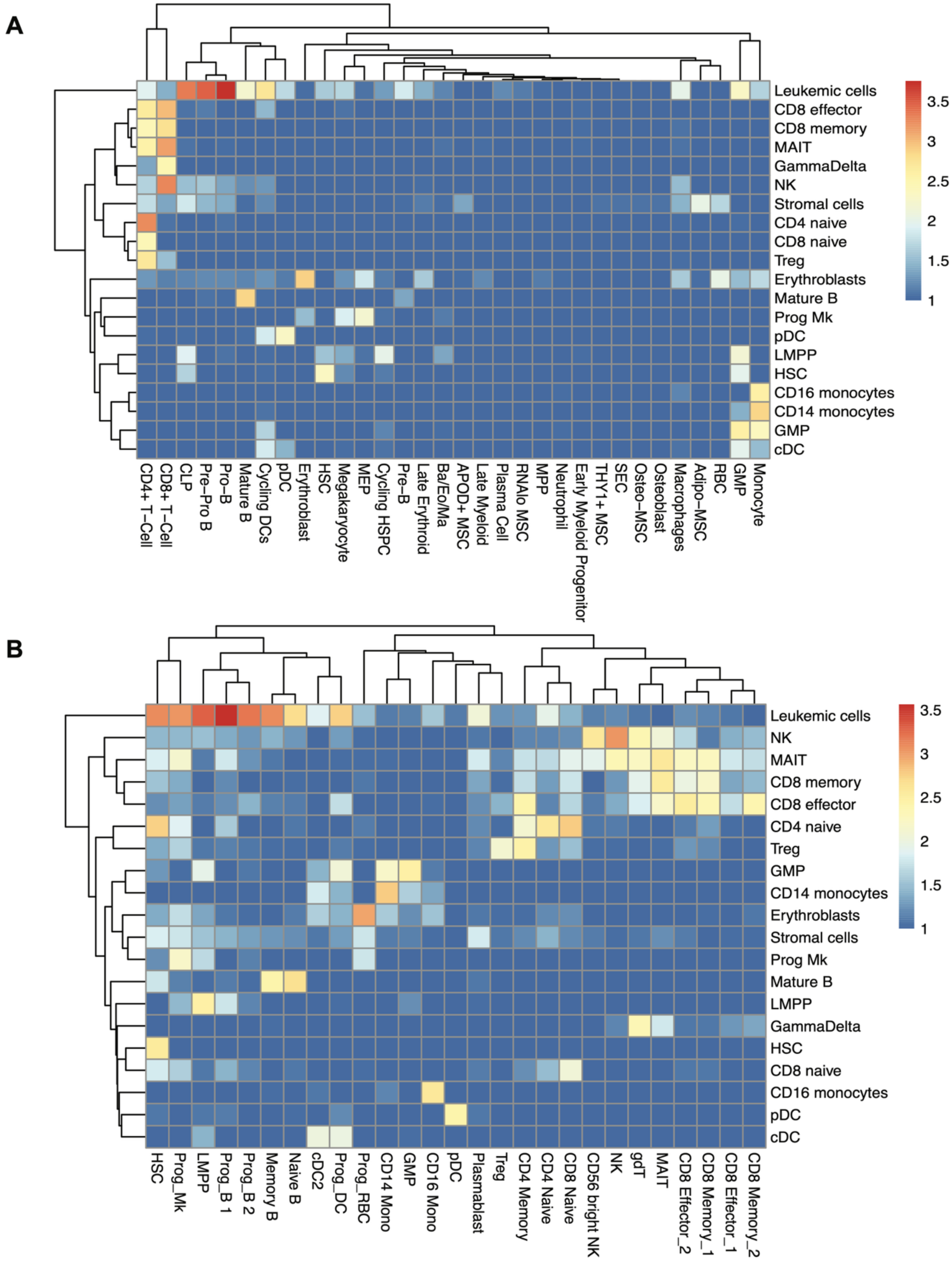


Supplementary Figure 3. Identification of cell types using SingleR. A) Heatmap representation of the scores assigned to the cell clusters using the annotation of Bandyopadhyay et al, Cell 2024. B) Heatmap representation of the scores assigned to the cell clusters using the annotation of Stuart, Butler et al, Cell 2019. For both, A and B, in the y-axis the name of the cell clusters assigned in this study, while in the x-axis the clusters identified in the respective publications.

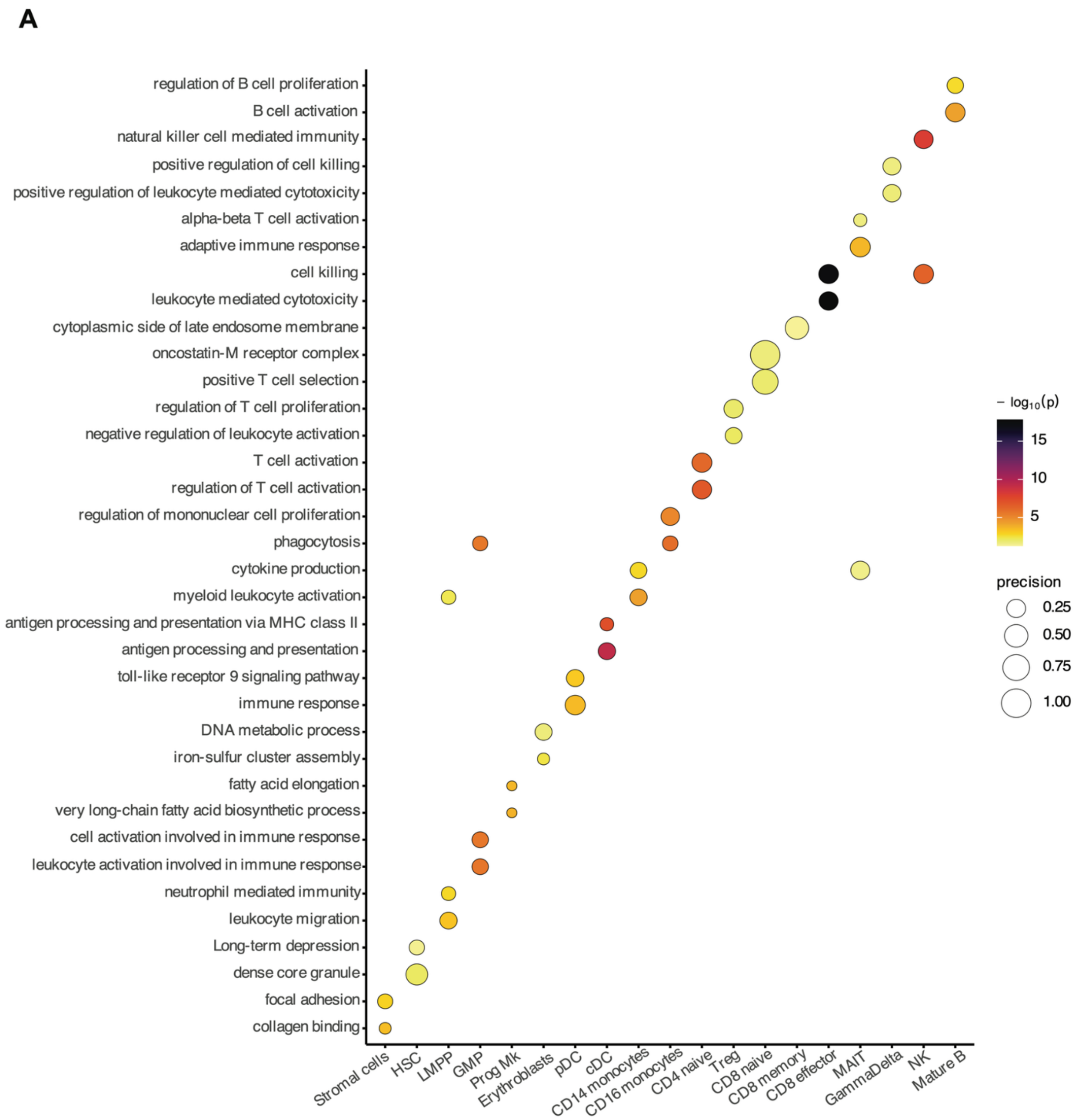


Supplementary Figure 4. Dot plot showing enriched gene ontology (GO) terms of differentially expressed genes in the cell clusters.

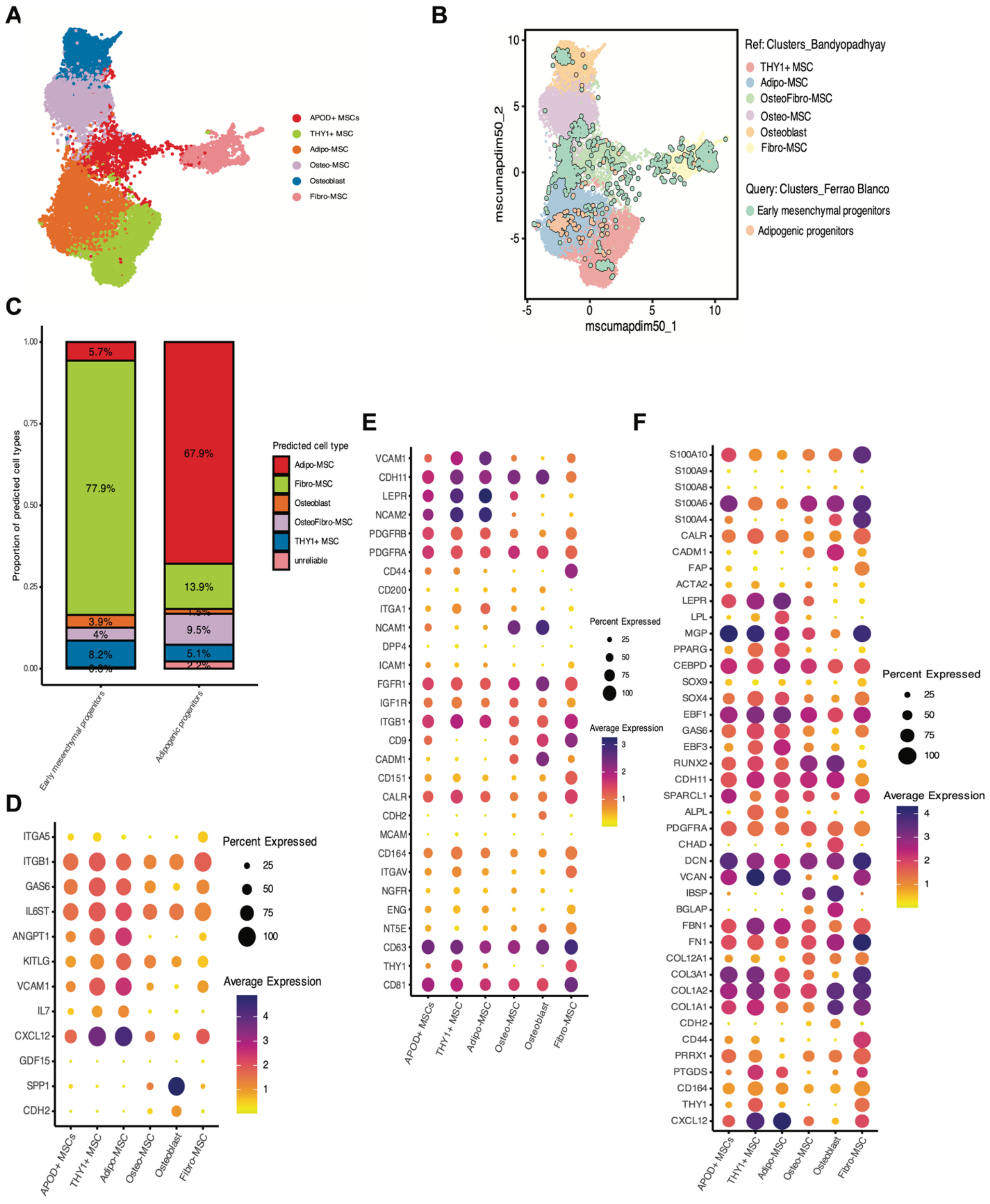


Supplementary Figure 5. Comparison of the stromal populations identified in this study with a human bone marrow stromal cell atlas used as a reference, Bandyopadhyay et al, Cell 2024. A) UMAP representation of the stromal clusters identified in the reference dataset. B) UMAP representation of the scRNAseq data of the two stromal populations identified in this study projected in the reference dataset. C) Barplot showing the reference-based annotation of the stromal populations identified in this study. D) Dotplot representation of the gene expression of widely studied niche factors in the reference dataset. The size of the dot indicates the percentage of cells expressing the gene and the color indicates the average expression level. The expression of these genes in our dataset is shown in Figure 5A. E) Dotplot representation of selected cell surface markers in the reference dataset. The size of the dot indicates the percentage of cells expressing the gene and the color indicates the average expression level. The expression of these genes in our dataset is shown in Figure 4A. F) Dot plot representing the gene expression of selected markers in adipogenic and early mesenchymal progenitors on the reference dataset. The expression of these genes in our dataset is shown in Figure 2D.

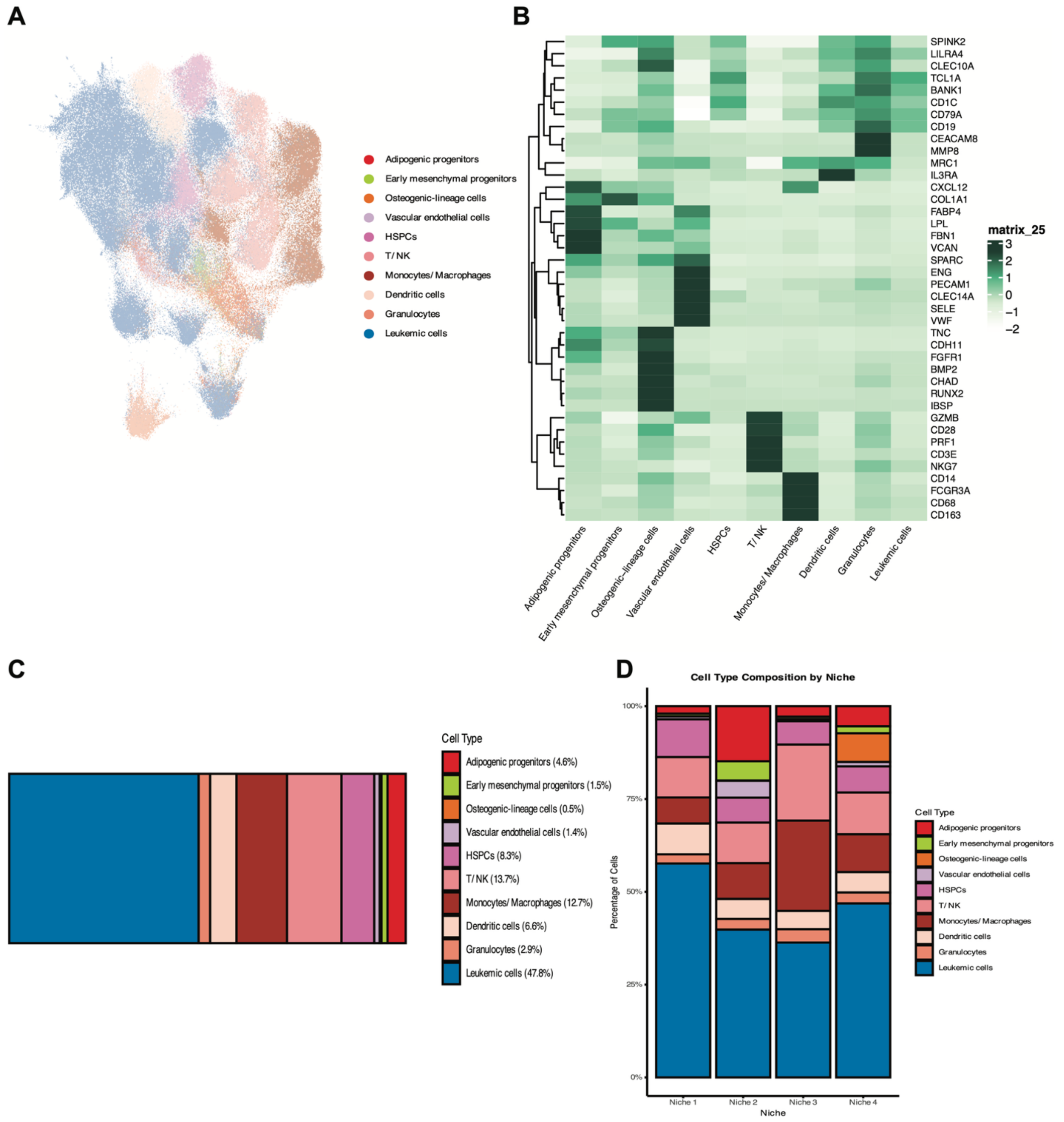


Supplementary Figure 6. Spatial transcriptomic analysis on the B-ALL microenvironment. A) Uniform Manifold Approximation and Projection (UMAP) plot showing the cell clusters identified in the Xenium data. B) Heatmap showing the average gene expression per cluster of selected top markers. C) Barplot representing the percentage of each cell type in the dataset. D) Barplot showing the cell type distribution between the four niches.

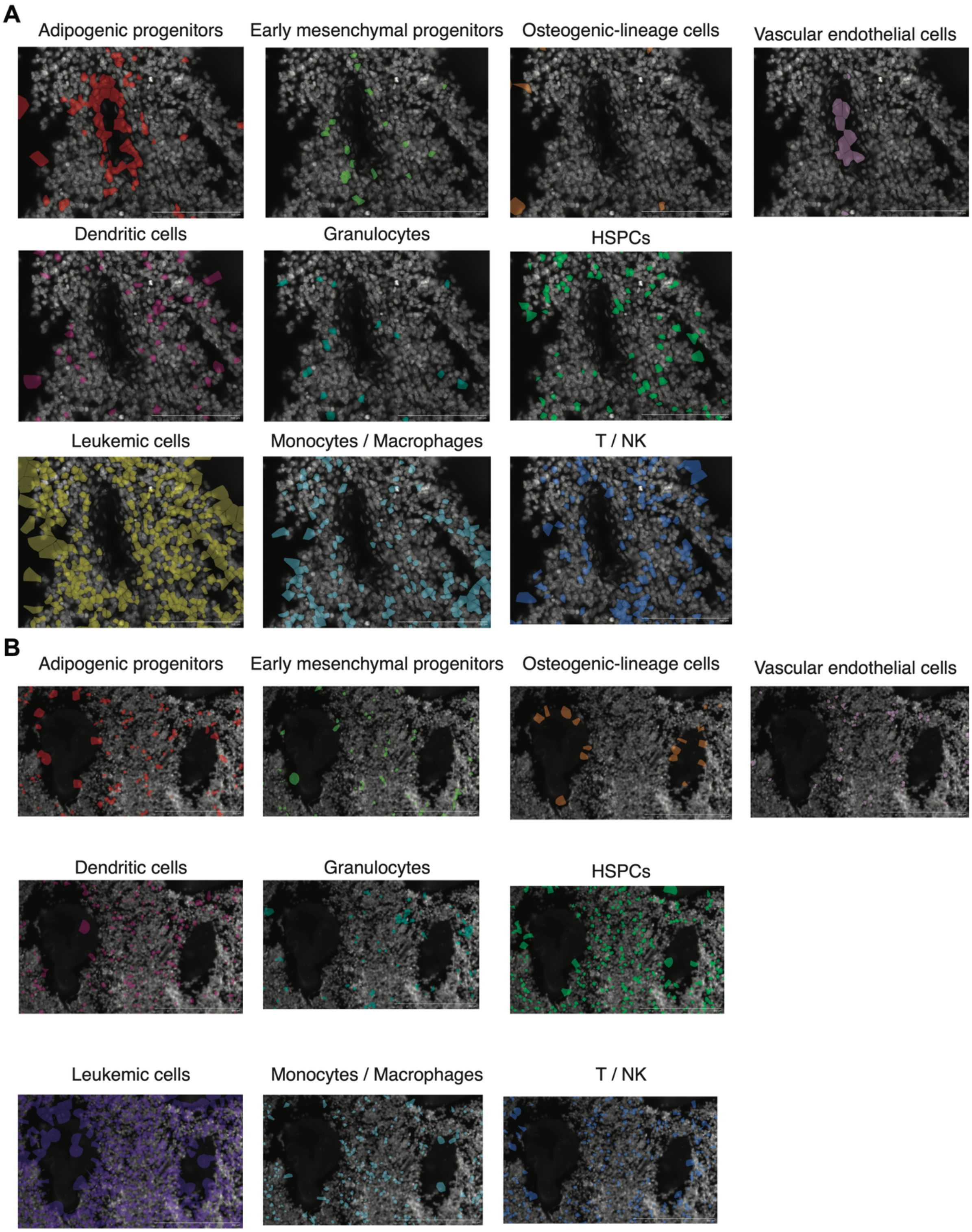


Supplementary Figure 7. Cell type localization in the selected regions, as in Figure 3D (A) and as Figure 3E (B).

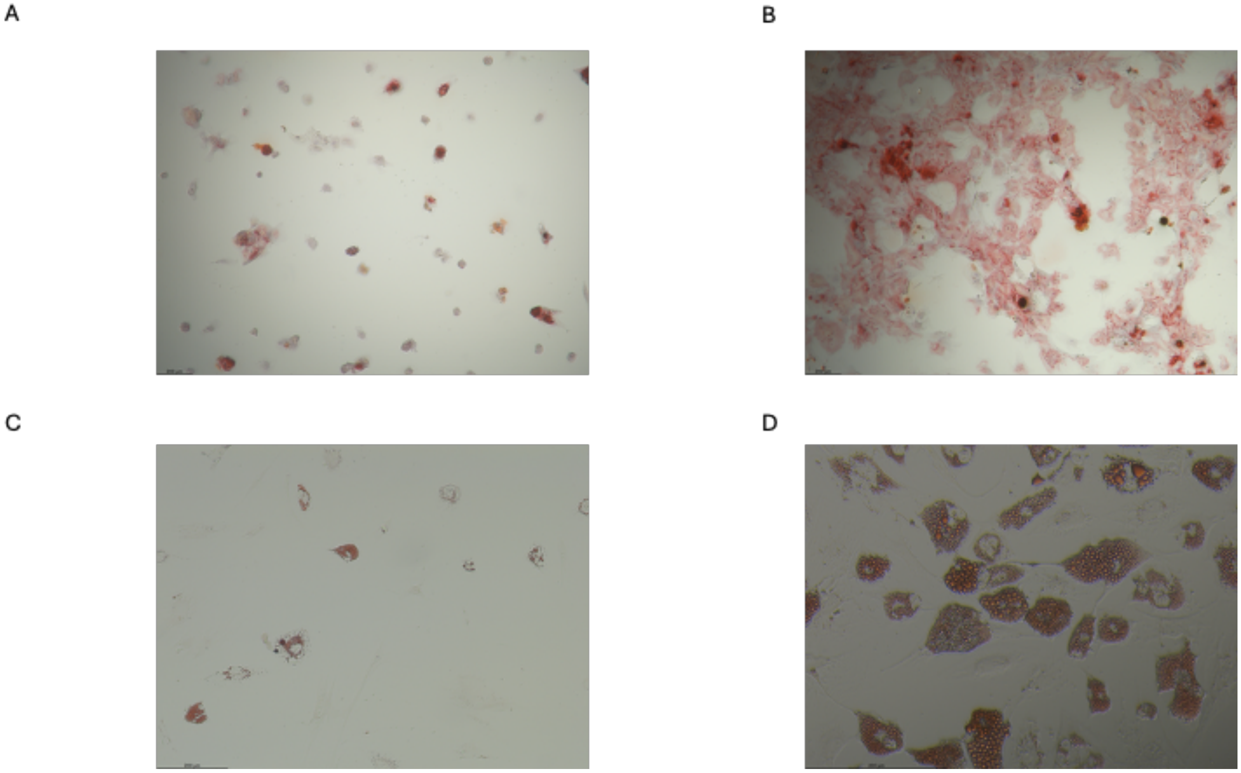


Supplementatry Figure 8. Adipogenic and osteogenic differentiation assay in sorted stromal populations, early mesenchymal populations (A, C) and adipogenic progenitors (B, D). Oil red O (C, D) and Alzarin red (A, B) are used to stain oil droplets and calcium deposition, respectively.

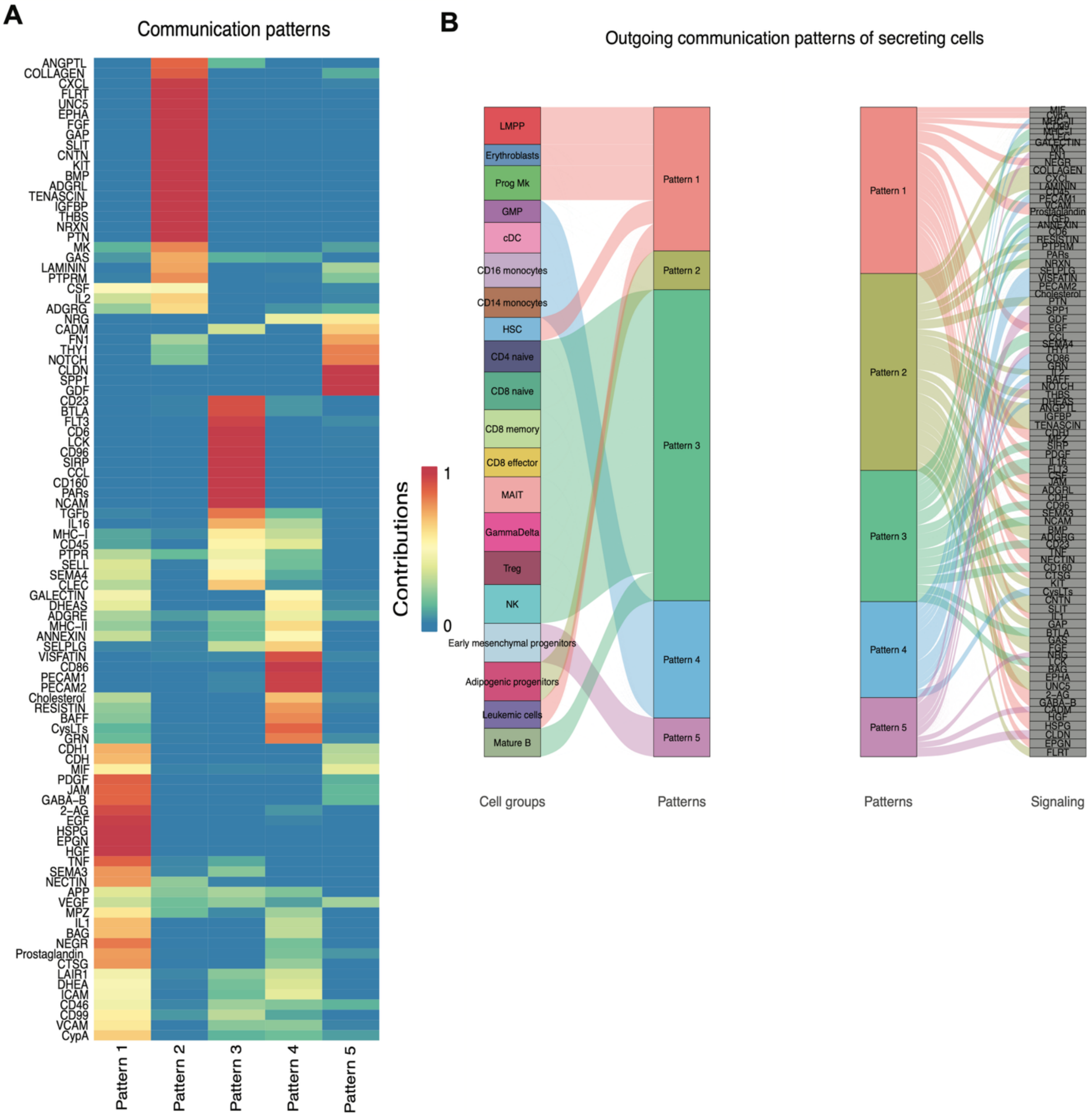


Supplementary Figure 9. Analysis of the communication network in the B-ALL niche. A) Non-negative matrix factorization (NMF) to identify modules in the cell-cell interaction pathways between niche cells and leukemic cells. Heatmap depicting the cell-cell communication pathways contributing to each pattern. B) Clusters of cells are grouped into each module.

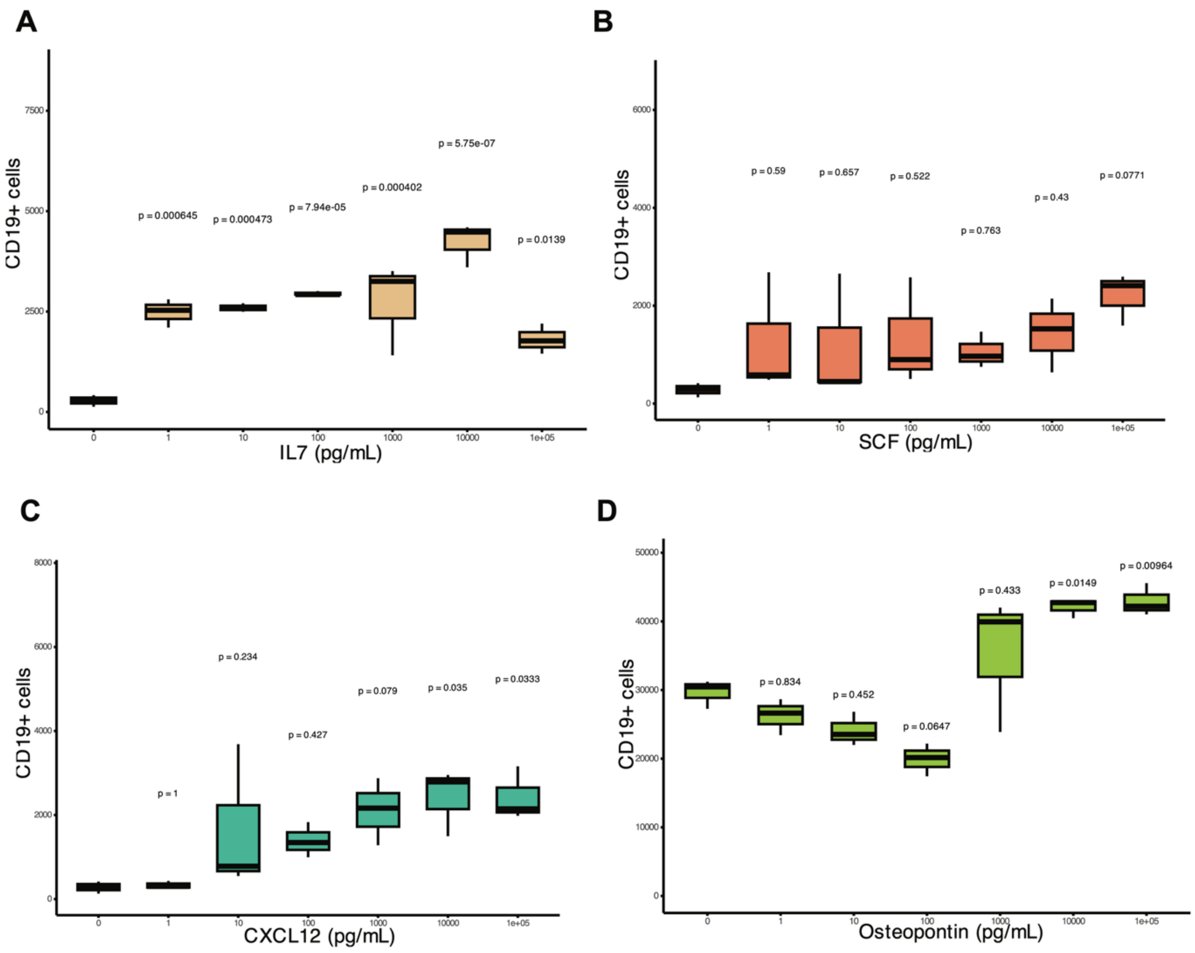


Supplementary Figure 10. Cytokine-mediated support to B-ALL cells ex vivo. Absolute number of CD19+ leukemic cells per well after seven days of monoculture with different concentrations of interleukin 7, SCF, CXCL12 and Osteopontin. Data is presented as mean

± standard deviation. Statistical significance was assessed by a **one-way ANOVA** followed by **Dunnett's post-hoc test** to compare each cytokine-treated condition against the untreated control (0 ng/mL).

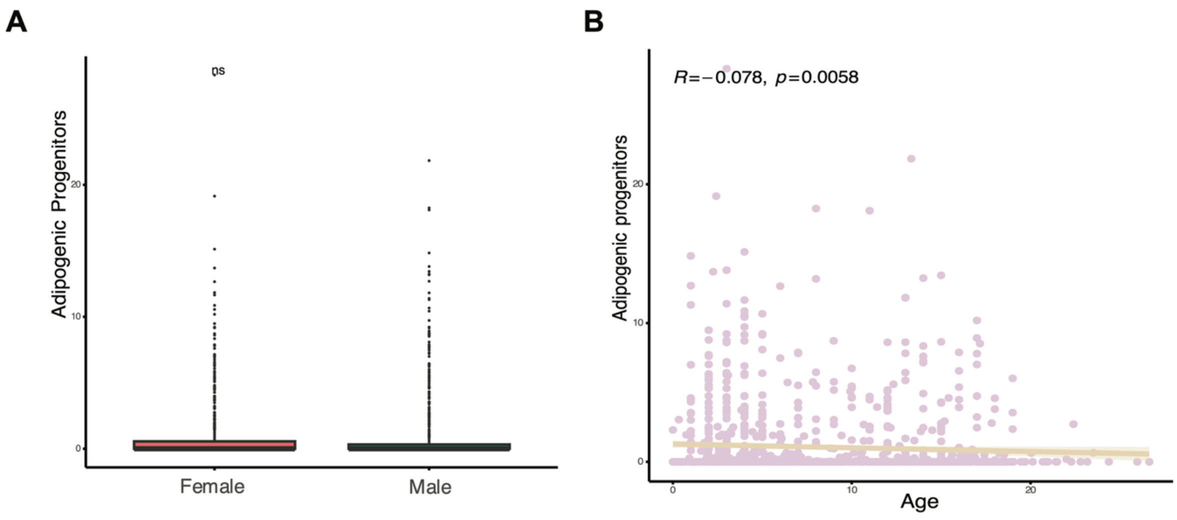


Supplementary Figure 11. Analysis of Cell type deconvolution. A) Comparison of the adipogenic progenitors between male and female patients. B) Correlation analysis between adipogenic progenitor signature score and patient age in bone marrow samples (spearman correlation).
